# Tropical forest dynamics correspond to fair games in economic theory of financial markets

**DOI:** 10.1101/2021.11.09.467956

**Authors:** Yue Lin, James Rosindell, Uta Berger, Helge Bruelheide, Jens Kattge, Volker Grimm

## Abstract

Ecological and economic systems both comprise of autonomous adaptive agents. It is thus possible that similar mechanisms determine the organization of both these complex systems. Indeed several economic theories have already been successfully applied in an ecological context. Here we show that ‘efficient market theory’ in economics, where future earnings are distributed between competitors by a ‘fair game’, corresponds to fitness-equalizing mechanisms of coexistence in ecology. In contrast to stabilizing mechanisms, which promote coexistence by giving each species an equilibrium abundance that is resilient to perturbations, equalizing mechanisms promote coexistence without such resilience by minimizing the net fitness differences between species. However, identifying stabilizing and equalizing mechanisms from the short time-series data that are typically available in ecology is challenging. We used techniques from economics that are applied to collections of short time-series from a system. We found that observed species abundance dynamics in a neotropical forest are generally in agreement with efficient market theory implying a dominant role of equalizing mechanisms, which finding quantifies and supports what was generally believed about that specific forest system. Our study highlights that complex systems from ecology and economics share common features suggesting the possibility of further synergy between ecology and economics in future.

## Introduction

Both ecological and economic systems comprise of competing autonomous individuals or agents each pursuing a certain purpose (Grimm et al. 2005). They are thus likely to share fundamental properties and have common ground in analytical techniques (Nekola and Brown 2007; May et al. 2008). Nevertheless, ecology and economics have developed along disjunctive pathways, leading to seemingly different concepts, theories, and methodologies. Some beneficial concepts and approaches developed in each discipline might therefore be usefully transferred to the other and help overcome limitations in both (Hammerstein and Hagen 2005). For example, the law of supply and demand for the theory of island biogeography (MacArthur and Wilson 1967), evolutionary game theory (Smith 1982), stability of ecological and financial networks (May et al. 2008), and Pareto optimality for genotype space (Shoval et al. 2012).

The efficient market theory (Fama 1970; Barnett and Serletis 2000; Poitras 2010) is a corner stone in financial economics, which states it is impossible for investors to “beat the market” because all available information is already incorporated and reflected into prices due to market efficiency. Here we show links between efficient market theory and ecological coexistence theory, using concepts and methods from the efficient market theory to explore mechanisms underlying coexistence in a species-rich tropical forest.

From the perspective of modern coexistence theory, ecologists have recognized two broad types of mechanism for explaining community assembly (Chesson 2000; Adler et al. 2007; HilleRisLambers et al. 2012): stabilizing mechanisms that generate negative density-dependence for promoting coexistence among different species, and equalizing mechanisms that reduce overall fitness differences among species with stochastic processes typically playing an important role. Both stabilizing and equalizing mechanisms can operate simultaneously, and their relative importance may vary among different communities (Chesson 2000; Gravel et al. 2006). The relative importance of stabilizing and equalizing mechanisms cannot, however, be fully evaluated simply by looking at static patterns such as species abundance distributions (Adler et al. 2007; Zhang et al. 2009). Analyses of dynamic patterns are more appropriate but are scarcely used in ecology (Keil et al. 2010; Rosindell et al. 2012) due to the lack of long empirical time series (Magurran et al. 2010).

Understanding how large numbers of species can coexist represents a significant challenge in ecology. A major limitation to understanding is the lack of empirical tests on co-occurring species dynamics to infer, via time-series analysis, the underlying mechanisms that lead to coexistence (Clark and McLachlan 2003; Rosindell et al. 2012). For most ecological communities, such temporal tests and analyses are limited by the demands for long census time-series, which are rarely collected in the field. Large numbers of short time-series are more readily accessible, but current techniques used in ecological research are not well suited to dealing with such data. These factors have made it difficult to reveal the relative importance of different mechanisms underlying species coexistence.

Here we adopt concepts and methods from economics to show that even short-term time series contain relevant information for the purpose. This is possible since the dynamics of species abundance under ecological equalizing mechanisms are analogous to the financial dynamics asserted by ‘efficient market theory’ (Fama 1970; Barnett and Serletis 2000; Poitras 2010). Both dynamics correspond to the ‘martingale’, a stochastic process with certain properties that emerge from ‘fair game’ dynamics where no party has a fundamental advantage (Figure 1). In contrast, under strong stabilizing mechanisms, species abundance dynamics become stationary processes where species abundances are regulated towards a target level. We also show how the same concepts and methods can, under different transformations, relate to ecological neutral theory (Hubbell 2001) with and without dispersal limitation.

**Figure 1:**
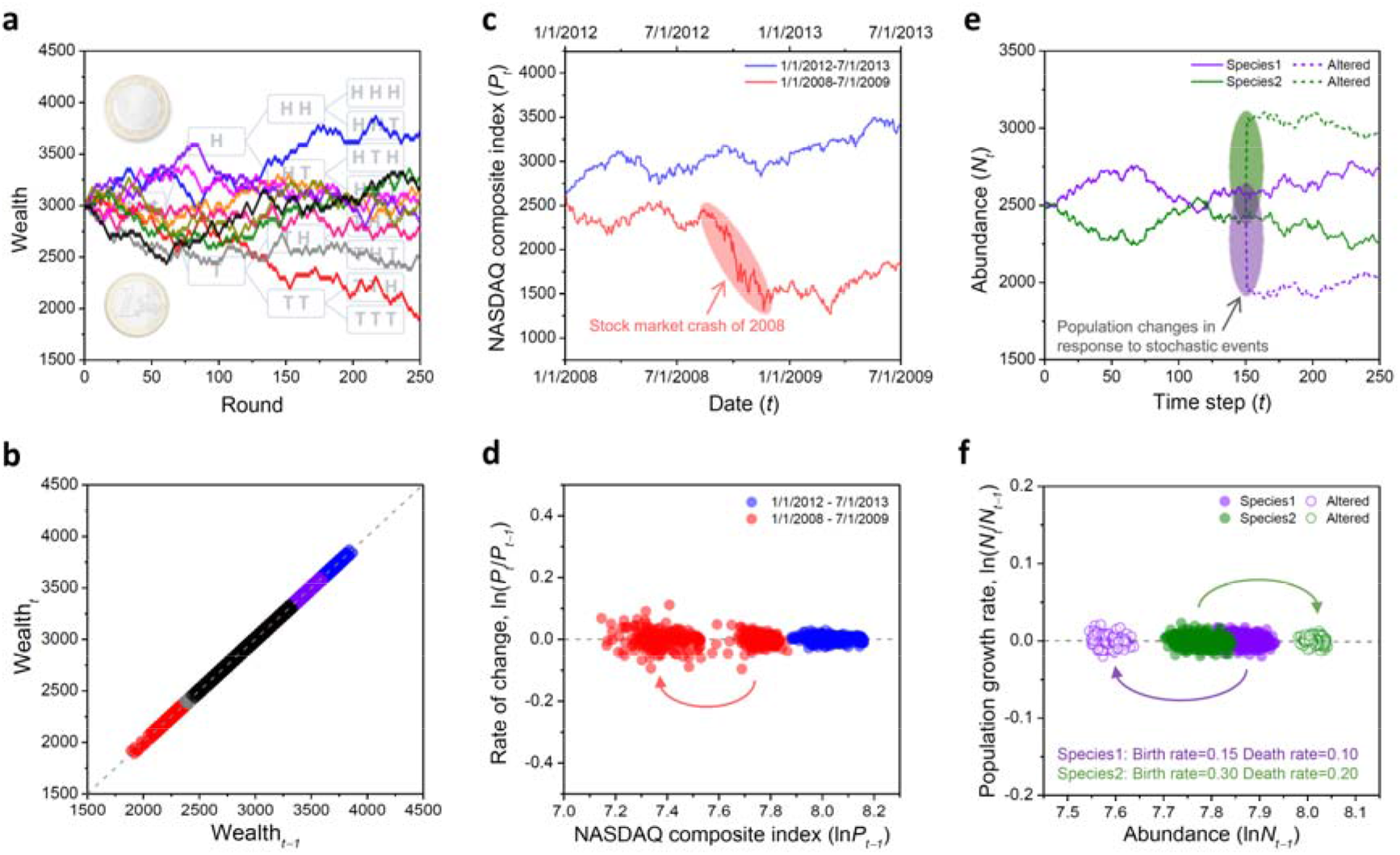
Examples of fair game dynamics (martingale processes). (*a*) A simulation of fair coin-tossing (10 replicates) where players gain or lose 30 arbitrary units in each round. (*b*) The expected wealth at time *t* as a function of the previous wealth at time *t*−1 using the time-series from panel a). (*c*) Historical data of the NASDAQ composite index (data from Google finance). (*d*) According to efficient market theory, the expected stock change rate is zero and is independent of past stock prices. The red time-series shows a sudden crash (see arrows in panel c and d), after this crash the stock price does not show a trend of return to previous levels (martingale behaviour). (*e*) An individual-based model simulation of two species dynamics based on equalizing mechanism of demographic trade-offs. (*f*) Under equalizing mechanisms, the expected population growth rate is zero and is independent of species’ abundance, implying no frequency-dependent regulations. This property becomes visible after a randomly chosen half of trees die perturbing the system: although species 2 suddenly exceeds Species 1 due to its higher absolute birth rate, but when they reach the equilibrium carrying capacity, neither species returns to pre-disturbance levels.

We evaluated multiple time-series of co-occurring tree species from a species-rich tropical forest by using econometric techniques known as ‘panel unit root tests’ (Hsiao 2003, 2007). These techniques were originally developed in economics for analyzing survey data (known as ‘panel data’ to economists) consisting of many short but associated time-series, often more than twenty time-series each with only five or six time points.

Here we used a data set from a 50 ha-plot within a species-rich tropical forest on Barro Colorado Island (BCI), Panama (Hubbell 2001; Condit 1998; Condit et al. 1999, 2012). The data comprise the abundances for more than 300 species and span 29 years with 7 census time points (figure 2). This data set has been widely used and thoroughly analyzed with various techniques before and the specific conditions of this neotropical forest are well-known (Condit et al. 2012; Chisholm et al. 2014; Kalyuzhny et al. 2014, 2015). We are using it mainly to illustrate how panel unit root tests from econometrics can be used to analyze short-term time-series of ecological data and to identify mechanisms. In addition, we explored the correlations between functional traits and demographic rates in BCI forest to provide an initial understanding of how fitness-equalizing mechanisms and ‘fair game’ dynamics emerge. The main aims of our study are to further the useful analogy between ecological and economic systems, and to demonstrate new techniques for detecting the processes of species coexistence.

**Figure 2:**
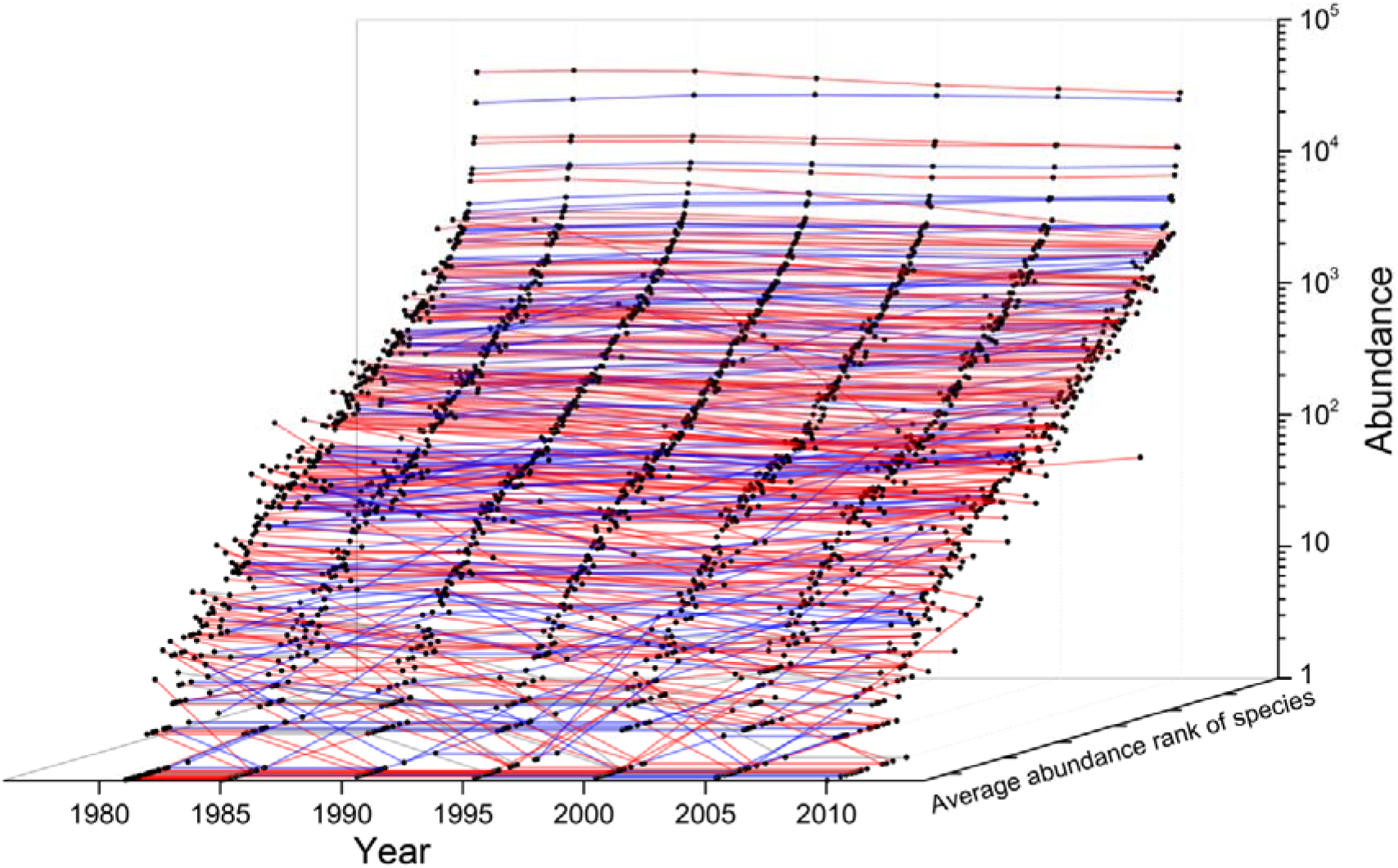
Abundance of 322 tree species (DBH ≥ 1 cm) over 29 years in a 50 ha-plot of species-rich tropical forest on Barro Colorado Island. Species were ranked in order of their average abundance across the entire time-series. Red lines indicate species whose abundance decreased, blue lines indicate species whose abundance increased, and gray lines indicate species whose abundance remained unchanged over the entire period of censuses.

## Theory

### Martingale Processes and Fair Games

A ‘martingale’ is a stochastic process, the net outcome of successive rounds of a ‘fair game’ where the next expected value is equal to the previous value regardless of past information. By definition it has the property

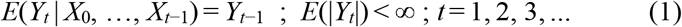

where *Y_t_* represents wealth in the game at time *t* and *E*(*Y_t_*) gives the expected value of *Y_t_* given the system’s condition at all previous times: *X_t_* for *t* = 0, 1, …, *t*−1 (Barnett and Serletis 2000). The key feature of this definition is that it is impossible to use any of the past history of the game to develop a winning strategy where *E*(*Y_t_*) > *E*(*Y*_*t*−1_). One can equivalently write that

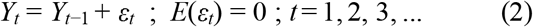

Where *ε*_*t*_ is the so called ‘martingale difference’, the outcome of a round of the game at time *t*, which is a random term with zero mean and independent distribution (e.g., white noise).

Consider a simple fair game of coin-tossing: the probability of getting a head or a tail is always 1/2 and the player respectively gains or loses the same amount of money. The expected profit of each round, the martingale difference, is always zero, even if we know the full history of the game. If a player’s wealth in round *t*−1 is *Y*_*t*−1_, then the expected wealth after the next trial, *Y_t_*, remains *Y*_*t*−1_ and the time-series of player’s wealth is a martingale even if trajectories sometimes appear to follow deterministic trends (Figure 1a, b).

### Efficient Market Theory

The financial economic model proposed by efficient market theory is a martingale on a logarithmic scale. One can formally define this as

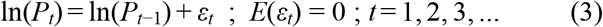

where *P_t_* is the price (of a stock for instance) at time *t* and *ε*_*t*_ is the martingale difference.

Equation (3) can be obtained from equation (2) by setting ln(*P_t_*) = *Y_t_*; the equation implies that it is impossible to make economic profits based on historic information in the market (Figure 1c, d). In economic theory it makes intuitive sense to work on a logarithmic scale because profits are typically reinvested and so the quantity of gain or loss in each round is a proportion of the total assets. For example, a fair game of coin tossing like the one described above, but where the amount gained or lost in each round is a percentage of the player’s total wealth instead of a fixed quantity. Equation (3) describes a non-stationary process giving prices that vary over time. However, the difference between successive logged prices (the rate of change on a logarithmic scale) is stationary and consists of values drawn from a time independent, normal distribution with zero mean. The typical method for testing whether a time-series of prices *P_t_* was created by a martingale under efficient market theory is based on a generalization of equation (3) given by

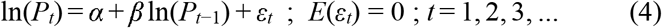

where *ε*_*t*_ is the random term of martingale differences usually assumed as white noise (Barnett and Serletis 2000; Poitras 2010), and *α* and *β* are fitted constants. If *α* = 0 and *β* = 1 then equation (4) reduces to equation (3) producing a martingale. The tests used, known as ‘unit root’ tests in the econometrics literature (Barnett and Serletis 2000; Poitras 2010), are therefore focused on determining if *α* is significantly different from 0 and *β* significantly different from 1.

### Ecological Martingales

Instead of a stock (or business) with assets and a price, now consider a species with individuals of a certain abundance. In this context the principles of efficient market theory have an analogy in ecology and the same tests and equations can be applied. In an ecological setting, efficient market theory would mean that no species can evolve or adapt a set of traits based on past environments that would guarantee success (increases in abundance) in future environments. This implies a purely ‘equalizing’ mechanism (Chesson 2000) because where differences between species exist, they will be irrelevant in determining future success. There are no stabilizing mechanisms involved here because there is no feedback between past and future failures and successes; the population growth rate for all species is thus unpredictable and independent of past abundances. In such a system, the abundances of species do not necessarily return to their previous levels after perturbations such as environmental disturbances (Figure 1e, f). This can be formalized by replacing the price of a stock *P_t_* with the abundance of a species *N_t_* in equations (3, 4) so that for our statistical tests we are using

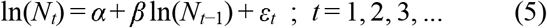

where *ε*_*t*_ is a random term representing demographic and environmental stochasticity as independent draws from a normal distribution with zero mean and variance 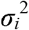 (white noise). Equation (5) in fact happens to be identical to the linear form of the stochastic Gompertz model, which is frequently used in ecological studies for modelling population dynamics (Ives et al. 2003).

There are two noteworthy distinctions between the economic and ecological martingales. Firstly, in economics the independence of future success from *all* former states of the system is critical to being a martingale, as traders or businesses will look at historic information about a stock to make decisions, but no one can beat the market because all information has been already priced into the market as suggested by efficient market theory. In ecology, however, the independence of the current state of the ecosystem is the only important issue for the future success; this is because in ecology the past is only relevant where it has left physical relicts (such as in a seed bank). All previous settings are “stored” in the current state of the ecosystem (Grimm and Wissel 2004). Secondly, in ecology the agents in the system are individual organisms and the population (species abundance or price in our analogy) is an emergent result of the individuals and their interactions. Conversely, in economics the decision making agents are distinct from the quantity of interest (currency). For an individual agent to perform in ecology means creating offspring, whereas for an agent to perform in economics means creating more wealth.

In our ecological analogy of efficient market theory, population growth or decline is multiplicative and thus the time-series of logarithmic abundance for a species is a martingale as in equation (5). This arises where conspecific individuals all have very similar prospects of reproduction or death at any particular time, for example because of environmental fluctuations acting simultaneously on all conspecific individuals (Chisholm et al. 2014; Kalyuzhny et al. 2014; Jabot and Lohier 2016). Such fluctuations over short timescales must still average out over longer time periods so that there is no net growth (or decline) of the species on logarithmic axes; this is a condition of being martingale and is necessary to prevent extinction or unbounded growth.

In an alternative ecological scenario, each individual organism *independently* reproduces or dies (regardless of what other individuals are doing) and on average has a net lifetime reproductive success of 1, again to prevent extinction or unbounded growth. This scenario would arise where the greatest drivers of success are factors affecting individuals independently and it produces a martingale on *arithmetic* axes evaluated using a parallel to equation (5) for arithmetic space:

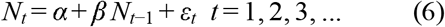

The change to arithmetic axes comes about because the rate of change in species abundance at a point in time is given by summing independent draws from a distribution of individual reproductive successes, instead of by drawing once from such a distribution and applying that same result to all conspecific individuals as in equation (5). Martingales on arithmetic axes can be analyzed with the same ‘unit root’ tests, but are less germane in economics because units of currency are not independent agents, so price changes will tend to only be multiplicative with strategies producing profit or loss as a percentage of the total amount invested.

To extend the species dynamic model of equation (5) to a community scenario with many species, we add an index *i* obtaining:

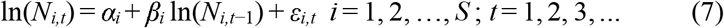

where *N_i,t_* is the number of individuals in species *i* at time *t*. Similarly, in arithmetic space, we can extended equation (6) with a species index *i* to give

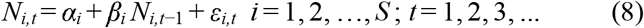

These economics inspired methods help to detect stabilizing and equalizing mechanisms separately in ecological systems. Particularly, equation (7) in logarithmic space represents dynamics under fluctuations of environmental stochasticity, whereas equation (8) of arithmetic space emphasizes demographic stochasticity. Furthermore, we have shown that equations (7) and (8) can, under different transformations, relate directly to ecological neutral theory (Hubbell 2001) including nearly neutral models of ecological trade-offs (Lin et al. 2009), and the more general concept of symmetric models (see Appendix A for details).

## Material and Methods

To test for the presence or absence of martingale processes underlying the multiple time-series of species abundance, we applied ‘panel unit root tests’. The basic concept of these tests is similar to that of meta-analysis: it combines the statistical information of all the time-series across different sections (species in our case) to draw an overall conclusion (Maddala and Wu 1999; Im *et al*. 2003). The general model of panel unit root tests applied to co-occurring species is shown in equation (7) and can be rewritten as:

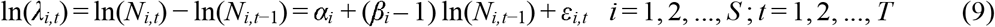

where *λ*_*i,t*_ is the finite population growth rate of species *i* at time *t*, *α*_*i*_ and *β*_*i*_ are species-specific constants, *ε*_*i,t*_ is a random term (i.e., demographic and environmental stochasticity) as before with mean zero and variance 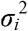. We need to test whether *α*_*i*_ = 0 and *β*_*i*_ = 1 for all *i*, which would make the system equivalent to a martingale model representing ‘fair game’ dynamics and equalizing mechanisms (Chesson 2000). If *α*_*i*_ ≠ 0 and |*β*_*i*_| < 1, the dynamics of ln(*N_i,t_*) approach a time-independent stationary distribution with a mean of *α*_*i*_/(1−*β*_*i*_) and a variance of 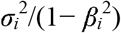 implying stabilizing mechanisms of negative frequency-dependent regulation (Ives et al. 2003). In order to test for martingales in arithmetic space, rather than logarithmic space we simply performed all the same procedures on *exp*(*N_i,t_*) as well as on *N_i,t_* itself.

For completeness, we used three different panel unit root tests that are flexible and widely used: ADF-Fisher χ^2^ test (Maddala and Wu 1999), IPS test (Im et al. 2003), and ADF-Choi Z test (Choi 2001) (see Appendix B for details, available online). The null hypothesis used for all these tests was *H_0_*: *α*_*i*_ = 0 and *β*_*i*_ = 1, ∀*i* ∈ {1,2,3,…,*S*} which means that, all species’ time-series are martingales corresponding to equalizing mechanisms. This is a stronger assumption and therefore easier to reject than one in which only a proportion of species follow martingales. Our alternative hypothesis is therefore that some (not necessarily all) species are not described by martingales, which would indicate that their dynamics are stationary processes influenced by stabilizing mechanisms.

To supplement the panel data unit root tests, we used linear regressions to test whether *α* = 0 and *β* =1, for each species individually. We also applied the Ljung-Box test (Ljung and Box 1978), which is known to be robust and is widely used to test the null hypothesis of independence in individual time-series. Furthermore, we applied the Rank version of Von Neumann’s ratio (RVN) test to each time-series; this test is powerful for detecting randomness in small sample sizes (Bartels 1982). Population growth rates were calculated following the standard methods of the Center for Tropical Forest Science (Condit 1998; Condit et al. 1999).

We tested our model with the data set of the 50 ha-plot on BCI, Panama (Condit 1998; Condit et al. 1999) (Figure 2). The data set contains seven censuses (1981, 1985, 1990, 1995, 2000, 2005 and 2010), all free-standing trees in the plot with stem diameter ≥ 1 cm (at 1.3 m above the ground) were mapped, tagged, and identified to the species level. The data were collected according to standard methods of the CTFS studies (Condit 1998; Condit et al. 1999). Over the past three decades there has been high turnover on BCI, with more than half of the initial individuals being replaced; this time period corresponds to an entire generation for tropical forest trees (Figure S1, available online). This has caused large changes in species composition and thus provides an eligible data set for analyzing species dynamics of a forest community (see Appendix B for details, available online).

To evaluate the statistical power of the three panel unit root tests, we first generated time-series data sets by simulation with known parameters. Then we tested the generated data using each of the three methods. In our simulations, each generated panel data set consists of 322 species (time-series) of 7 time points, the same as the empirical data set from BCI. Each time series was generated by application of equation (7) to produce data on a log scale, and repeated separately using equation (8) to produce another data set on an arithmetic scale. In each data set the species were permitted to have different values of *α*_*i*_ and *β*_*i*_. The values of *β*_*i*_ were drawn from a normal distribution with a standard deviation of 0.02 and a mean of 0.1, 0.2, 0.3, 0.4, 0.5, 0.6, 0.7, 0.8, 0.9 or 0.99, corresponding to ten different scenarios of stabilization force strength. Thus different tests ranged from very strong stabilization forces (mean *β*_*i*_ = 0.1) to weak stabilization forces (mean *β*_*i*_ = 0.99). The *β*_*i*_ were truncated at 0.99 so that any values of *β*_*i*_ were set at 0.99 exactly, this was done because the stabilizing forces become too weak and break down as beta gets larger than this boundary. The 322 species in simulated data were cross referenced to the 322 species observed empirically and the species specific parameter *α*_*i*_ (for *1*≤*i*≤*322*) was chosen to match the long-term mean abundance of each simulated species to its empirical counterpart. Specifically, the empirical abundance of species i is given by 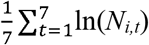 and the simulated log-term mean abundance, maintained by stabilizing forces, is given by *α*_*i*_/(1− *β*_*i*_). Note that the latter is undefined when *β*_*i*_ = 1 because in this case there are no stabilizing forces acting to maintain the abundance. Thus rearranging 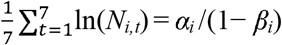 and taking the already chosen value of *β_i_* as well the empirical data *N_i,t_* enables us to solve for *α*_*i*_ and produce a dataset comparable to the empirical data but maintained by stabilizing forces of a given strength.

We generated 100 panel data sets at each stabilizing strength level by changing the seed of a pseudo-random number generator and performed panel unit root tests on each generated panel data set. The statistical power of a test is defined as the probability of rejecting the null hypothesis when it is indeed false, that is, the probability of not making Type II errors. The statistical power of the panel unit root test at a given stabilizing strength level is therefore estimated as the fraction of cases for which the null hypothesis is rejected (*P* < 0.05). Similar power calculations have previously been applied to ecological neutral theory in the context of detecting neutrality from species abundance distributions (Al Hammal et al. 2015).

In addition to simulated time-series datasets based on stabilizing forces, we also developed a simple individual-based tree competition model and applied panel unit root tests to this phenomenological model. In the tree competition model, 39 species coexist and compete for 10,000 empty grid-cells. Species dynamics can be determined by either stabilizing or equalizing mechanisms, providing an independent dataset with which to evaluate the performance of the panel unit root tests. See Appendix B for a more detailed description of the individual based model and Appendix C (available online) for the model code implemented in NetLogo 4.1.3 (Wilensky 1999).

To explore a potential explanation for the emergence of fitness-equalizing mechanisms and ‘fair game’ dynamics, we considered fitness-equalizing trade-offs based on species functional traits. Five functional traits were examined: 1) wood density, 2) seed dry mass, 3) leaf mass per unit area, 4) relative growth rate for saplings and 5) for large trees. We correlated these traits with mortality rates, recruitment rates and net population change rates for 134 tree species in the BCI plot where data were available (Wright et al. 2010) (see Appendix D for data and methodological details).

## Results

The panel unit root tests simultaneously included data for all co-occurring species in the BCI tropical forest community for which we had suitable time-series. In logarithmic space, all three tests failed to reject the null hypothesis (*P* = 1.000, Table 1), indicating that the population dynamics of tree species in the forest community are martingales, and are thus consistent with dominating equalizing mechanisms. The result remains robust when larger trees and saplings were tested separately in logarithmic scale (Table 1). We also applied the same three tests to species abundance time-series in arithmetic space (Table S1, available online), the results were partly consistent with what we found in logarithmic space, except that ADF-Fisher *χ*^2^ test rejected the null hypothesis for the test of all individual (*P* = 0.045, table S1) and for the test of only saplings with Diameter at Breast Height (DBH) ≤ 100mm (*P* = 0.038, table S1).

**Table 1.** Results of panel unit root tests for tree species abundance (logarithmic space) dynamics in a 50 ha-plot of species-rich tropical forest on Barro Colorado Island.

| | IPS test | ADF-Choi Z-statistics | ADF-Fisher $\chi^2$ |
| --- | --- | --- | --- |
|  | Statistic (Probability) | Statistic (Probability) | Statistic (Probability) |
| All individuals (DBH ≥ 1 cm) | 5.048 ($P = 1.000$) | 7.098 ($P = 1.000$) | 461.308 ($P = 1.000$) |
| Trees (DBH ≥ 10 cm) | 3.476 ($P = 1.000$) | 5.118 ($P = 1.000$) | 372.885 ($P = 0.970$) |
| Saplings (1 cm ≤ DBH < 10 cm) | 6.968 ($P = 1.000$) | 7.909 ($P = 1.000$) | 395.105 ($P = 1.000$) |

Null hypothesis: the existence of unit roots for all coexistence species; non-stationary dynamics (martingale processes) predicted by equalizing mechanisms. Alternative hypothesis: for some (but not necessarily all) species the unit root does not exist; stationary dynamics predicted by stabilizing mechanisms. Note it is not applicable to use the tests for all 322 species because some (38 species for all individuals, 113 species for larger trees, and 49 species for saplings) of the time-series are too short or incomplete (Appendix D, available online).

Single time-series analysis (Ljung-Box test and RVN test) on population growth rate of each species indicated that more than 96% of the time-series were temporally independent and random processes (Table S2, available online), which is consistent with the martingale property. The same result also holds true for both groups of larger trees and saplings (Table S2).

The average per capita growth rate for each species was consistently close to zero and did not depend on abundance (Figure 3), although the variation in average growth rate was larger for less abundant species, which is probably due to their small population sizes. Similar results have also been found in experimental microbial systems (Zhang et al. 2009). We found that none of the 74 species whose per capita growth rates were statistically different from zero showed a negative correlation between population growth rates and species abundances as predicted by stabilizing mechanisms (Figure S2, available online).

**Figure 3:**
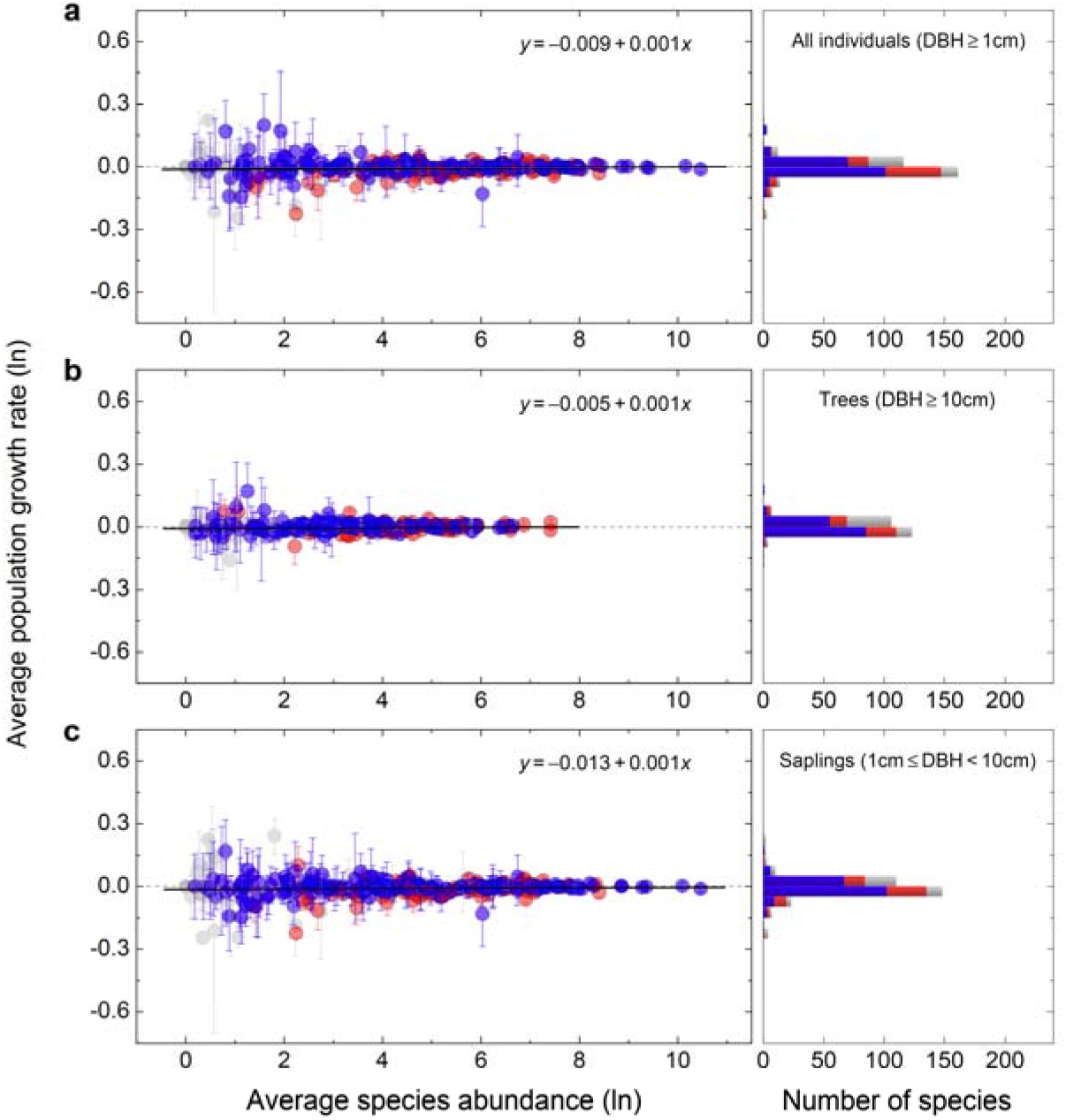
Log population growth rate (mean ± 1 SD) versus log average species abundance for all tree species in a 50 ha-plot of a species-rich tropical forest on Barro Colorado Island. Panels on the left show individual species as points, panels on the right show a histogram of species by population growth rate categories. (*a*) pooled data of all individuals (320 species, *r*^2^=0.001, *P* = 0.603). (*b*) data of all trees with DBH ≥ 10cm (243 species, *r*^2^=0.002, *P* = 0.519). (*c*) data of all saplings trees with 10cm ≥ DBH ≥ 1cm (310 species, *r*^2^=0.002, *P* = 0.423). The solid lines show fitted ordinary least-squares regression. The blue circles denote species whose growth rates did not differ significantly from ‘fair game’ dynamics of zero (*P* > 0.05; one-sample *t*-test). The red circles denote species whose growth rates differ significantly from zero i.e. they are driven by a significant component of stabilizing forces (*P* < 0.05; one-sample *t*-test). The gray circles denote species whose growth rates are not applicable for a *t*-test (*P* < 0.05; Shapiro-Wilks normality test). The horizontal bar charts on the right of each panel show totals of each of the three categories (fair game, stabilized, untested) across all abundaces. See Appendix D for further details.

We estimated the species-specific *α*_*i*_ and *β*_*i*_ parameters by fitting species abundance time-series data. In logarithmic space, more than 85% of species were consistent with the martingale property (*α*_*i*_ = 0 and *β*_*i*_ = 1), this also holds true when larger trees and saplings were separately estimated (Figure S3, available online). In arithmetic space, there were still more than 79% species consistent with the martingale property for all individuals, larger trees and saplings (Figure S4, available online).

We found strong correlations between demographic rates and all five functional traits (Figure 4). These correlations cancelled out each other in trade-offs to produce near constant net rates of population change across all species consistent with fitness-equalizing mechanisms and ‘fair game’ dynamics of co-occurring tree species on BCI.

**Figure 4:**
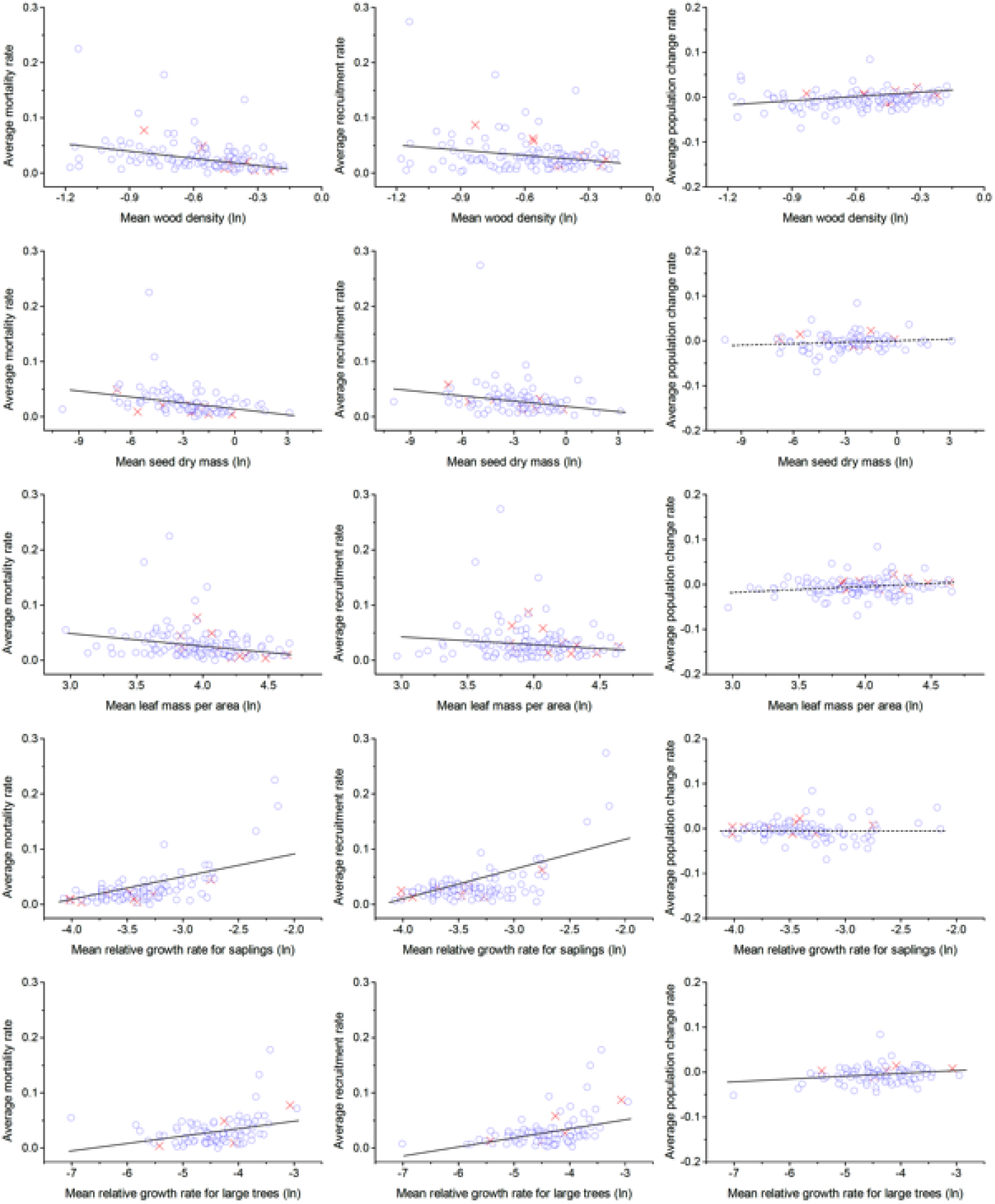
Trade-offs between 5 functional traits and demographic rates for 134 tree species in 50-ha plot on Barro Colorado Island, Panama. The blue circles denote species whose *α*_*i*_ and *β*_*i*_ are consistent with martingale properties (based on the results of all individuals showed in Fig. 5). The red crosses denote species whose *α*_*i*_ and *β*_*i*_ are not consistent with martingale properties. The lines show fitted ordinary least-squares regression. Solid regression lines were statistically significant (*P* < 0.05); dashed regression lines were statistically insignificant (*P* > 0.05). The five traits investigated were: mean wood density (gravity) (g cm^−3^), mean seed dry mass (g), mean leaf mass per area (g m^−2^), mean relative growth rate for saplings (cm cm^−1^ yr^−1^) and mean relative growth rate for large trees (yr^−1^). The regression for demographic rates and mean tree height were not statistically significant (*P* > 0.05) and are not shown.

The statistical power of three panel unit root tests used in our study is very high when stabilizing strength is strong or moderate, being naturally weaker when stabilizing strength is also weak (Figure 5). However, combining results of three tests (null hypothesis of ecological martingale is rejected if at least one test is statistically significant) strengthens the statistical power (Figure 5) even if stabilizing strength is very weak and time-series are very short (each time-series only contains 7 data points). The same result holds true in arithmetic space, but the statistical power is weaker than for logarithmic scale when stabilizing strength is very weak (see Figure S5, available online).

**Figure 5:**
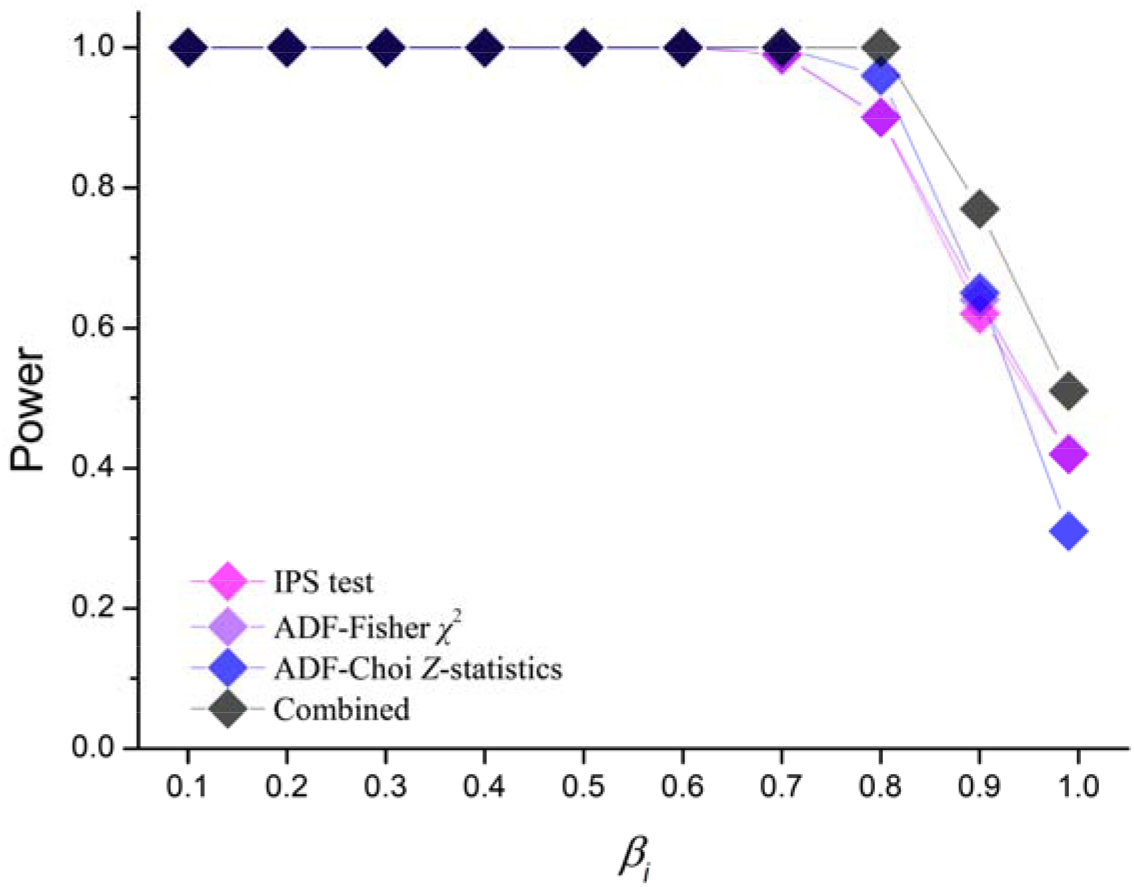
Statistical power of panel unit root tests on rejecting martingale null hypothesis as a function of the stabilizing processes strength ranging from 0.1 (very strong) to 0.99 (weak). Tests were performed in logarithmic space. The simulated panel data sets each contained 322 time-series of 7 data points and were generated based on the empirical data set of BCI. Just as is necessary with the empirical data set, we excluded a small number of the 322 time series from further tests as they were too short or incomplete (Appendix D, available online).

Our power tests based on simulation experiments of an individual-based tree competition model in which we imposed either stabilizing or equalizing mechanisms, showed that panel unit root tests correctly distinguish the processes in simulation data sets that only contain 39 species with 7 time points (Table S3 and Figure S6, available online). The empirical data from BCI forest that we study contains many more species (284 species for all individuals, 209 species for larger trees, and 273 species for saplings), than our simulation model data set, and so should be still more capable of detecting the alternative hypothesis of stabilizing processes (Hlouskova and Wagner 2006).

## Discussion

A significant difference between panel data tests and single time-series tests is in the hypotheses being tested: single time-series tests use a single martingale process as a null model and only apply to one species. In contrast, the panel tests we used here have a set of martingale processes as a null model, thus simultaneously considering all co-occurring species in the community, which is more relevant for addressing questions about whole community dynamics. In addition, panel data of the kind we used here contain more sample variability and more degrees of freedom compared to single time-series, so panel data analysis have a higher efficiency and accuracy of estimates (Hsiao 2007).

Our simulation experiment showed that the three panel unit root tests we employed are capable of distinguishing stabilizing processes from equalizing (martingale) processes based on short time-series (about 300 time-series with 7 time points), even when the strength of stabilizing processes is very weak (Figure 5, the mean value of *β*_*i*_ equals 0.9, for some species their *β*_*i*_ can reach 0.99). In addition, our individual-based model (Table S3) also demonstrated that panel unit root tests are very capable even when faced with data sets that are much less informative (only 39 species with 7 time points) than the data set of BCI forest (about 300 species with 7 time points) that we used here. In econometrics, studies of data generated by Monte Carlo simulations showed that panel unit root tests are often much more powerful than unit root tests of single time-series (Maddala and Wu 1999; Hsiao 2003; Im et al. 2003; Hsiao 2007), especially when panel data are very short but have large number of time-series (Hlouskova and Wagner 2006).

Intuitively one might assume that 29 years of 7 time points are too short to detect tree species dynamics and the mechanistic processes underlying. However, panel data analyses are specialized in compensating for the shortness of time-series by combining information from a large number of such time-series. Moreover, in the 50 ha tropical forest plot from which the data were taken more than half of the initially present individual trees were replaced over the past 29 years (Figure S1). This suggests a high amount of turnover and dynamical changes on BCI captured by the data set. Thus, our results obtained with panel data analysis do in fact reflect community dynamics rather than just small demographic fluctuations in a short period.

We find that even with extensive data and a powerful test, fair game dynamics corresponding to pure equalizing forces could not be rejected. The functional traits of species varied considerably and had an effect on demographic rates, but ultimately not on net population growth for each species (Figure 4), which remained constant again in line with the predictions of equalizing forces. Furthermore, for those few species showed non-zero growth rates, negative density dependence was not observed as would be expected from stabilizing forces (Figure S2). Together these findings suggest that stabilizing mechanisms do not play a dominant role in structuring the BCI forest community, which adds new and independent weight to conclusions from previous studies (Hubbell 2008; Condit et al. 2012; Kalyuzhny et al. 2014).

Equalizing and stabilizing mechanisms should lead to different kinds of species dynamics. Under ecological equivalence based on fitness-equalizing trade-offs, the species dynamics in the community would be a ‘fair game’ and thus a martingale process (Figure 1e, f). In ecological terms, these results imply that species abundance dynamics are not regulated by density-dependence (Ives et al. 2003; Dennis et al. 2006; Ziebarth et al. 2010; Knape and de Valpine 2012). Because negative density-dependence is a strong indicator of stabilizing niche differentiation (Adler et al. 2007; Levine and HilleRisLambers 2009; HilleRisLambers et al. 2012), species abundance dynamics that are governed by stabilizing mechanisms should approach stationary distributions with species-specific means and variances. The processes underlying such distributions do not have unit roots, differ between species, and thus are not martingales.

Our findings do not imply that stabilizing mechanisms are completely absent in the BCI plot, only that they may function together with the more important equalizing mechanisms and are not sufficiently strong to be detected from the data we had available. Many studies have shown that the stabilizing mechanisms stressed in niche theory are also important for maintaining species coexistence in ecological systems (Levine and HilleRisLambers 2009). It is noteworthy that there is a difference between stabilizing mechanisms that arise from traditional stabilizing forces (e.g., negative density-/frequency-dependence) and those that arise from other mechanisms that may be symmetric or neutral; for example in an ecological neutral model following Hubbell (2001) steady immigration from a fixed meta-community may act as a stabilizing force from the perspective of a local community (Chisholm et al. 2014). Other recent studies also illustrated the importance of spatial scale on the explanations of ecological patterns and processes (Chase and Knight 2013; Chase 2014), since both equalizing and stabilizing mechanisms are acting simultaneously at many scales (Chesson 2000; HilleRisLambers et al. 2012; Chase and Knight 2013; Chase 2014).

A martingale at first appears inconsistent with temporal autocorrelation in species abundances, which is often observed in empirical data. However, the presence or absence of such autocorrelation is also influenced by the timescales involved. For example, if we consider timescales shorter than the duration of fluctuations in environmental stochasticity, then we would expect successive points in the time-series showing consistent decline or growth in line with the environment at that time. If we consider successive points separated by a greater amount of time, then environmental conditions felt by the community at each time point could become de-correlated and changes in population size could be de-correlated likewise. These issues are quite separate from the interplay between stabilizing and equalizing forces, population size could revert to an equilibrium according to stabilizing forces, or fluctuate stochastically according to equalizing forces, and in either case do so with a degree of temporal autocorrelation at shorter timescales. The temporal scale of autocorrelation on BCI has been estimated to be 10 years (Kalyuzhny et al. 2015), which would only affect adjacent time points in the dataset with a 5-year census interval.

Future studies of the relative importance and contribution of stabilizing and equalizing mechanisms in community assembly and biodiversity maintenance should include different spatial and temporal scales (HilleRisLambers et al. 2012; Chase 2014). A promising next step would be to generate virtual data (Zurell et al. 2010) from community models representing different mechanisms and theories operating at a range of scales and check how well the econometric methods presented here can detect the relative contribution of the different mechanisms across scales. In this study, we focused only on the community dynamics operating at an ecological scale, when evaluating dynamics on an evolutionary scale in future work, the effects of mutation and speciation would also be important to consider, but are beyond the scope of our present work.

Recent studies suggest that environmental stochasticity plays a dominant role in driving community dynamics (Chisholm et al. 2014; Kalyuzhny et al. 2014; Jabot and Lohier 2016). Our results are consistent with those findings and suggest that ecological martingales also apply in such cases: no species can be ‘super-adaptive’ to the unpredictable events of environmental stochasticity in comparison with other species; species coexistence in an unpredictable stochastic environment is not stable, but indiscriminate and rather more like a ‘fair game’. We suggest that the role of environmental stochasticity in the causation of species responses (e.g., synchrony or asynchrony) could be evaluated in future studies by using econometric approaches related to those used here, for example the Granger causality test (Granger 1969).

Many have strived for a synthesis of theories to improve our understanding of biodiversity maintenance and functioning (Chesson 2000; Hubbell 2001; Chave 2004; Tilman 2004; Purves and Pacala 2005; Adler et al. 2007; Hubbell 2008; Allouche and Kadmon 2009; Lin et al. 2009; Zhang et al. 2009). We propose that these approaches could be combined with those used in economics to help achieve a still more inclusive synthesis by systematically comparing different ecological communities in various successional stages, under diverse environmental conditions. This would enable evaluation of the relative role and importance of different mechanisms during community assembly.

We have shown that species coexistence under fitness-equalizing stochastic events corresponds to a martingale (‘fair game’). However, the ecological martingale does not imply the absence of individual interactions and natural selection. In fact, individual interactions and natural selection are the driving forces for the emergence of ecological martingale. Here, we propose an analytical statement of our hypothesis of the ecological martingale: each individual’s life history strategy should keep abreast with the average performance of all other strategies in relation to demography. This idea is consistent with the Red Queen hypothesis (Van Valen 1973) and with the addition of weak selection into ecological neutral theory (O’Dwyer and Chisholm 2014; Rosindell et al. 2015). Similarly, this kind of interpretation on agent interactions and complex system dynamics was also made in economic systems for explaining the evolution of investment strategies and the emergence of arbitrage-free condition (Markose et al. 2005), which was known as the efficient market theory (Fama 1970), a corner stone of modern financial theory. Significantly, the debate on market efficiency in economy (Fama 1998; Shiller 2003) is very much like the debate on community assembly in ecology, which also implies a possibility of further synergy between the two fields in the future.

The mechanisms that assemble community is not completely neutral or niche, but somehow a continuous, between (Gravel et al. 2006), a more possible is dynamical nested structure, maybe not completely stable or equalizing, several niche in each niche can be one or more than one species. Those species within same niche are probably ecologically functionally similar (guild), that their dynamics are martingales. But within another niche, there is only occupied by one species at the moment and for this species the dynamic is stationary. In this case, the dynamics of functional guild is relatively stationary but species dynamics are still martingales, then maybe we should ask if community assembly functional traits or species? A rigorously test on this model could be applied.

Common patterns have been identified across different complex dynamical systems, such as physical, geological, cultural, economic and ecological systems (Grimm et al. 2005; Nekola and Brown 2007), so the same set of general mechanisms might indeed generate these patterns. Notably, the species abundance distribution empirically observed in ecology is very similar to the Pareto wealth distribution (also known as Pareto power law) found in economics (Nekola and Brown 2007; Scheffer et al. 2017), and the Pareto wealth distribution only occurs when the market is efficient, corresponding to a fair game dynamics (Levy 2006; Klass et al. 2007). In a recent study, Scheffer et al. (2017) found the inequality (Pareto distribution) in nature and society can be explained by stochastic events alone, this inequality can emerge naturally if wealth (or abundance) is subject to random losses or gains, even though no agent is superior to others. While our study signifies the link and common mechanism between the two systems by adopting models and techniques developed in economics for testing ecological hypotheses of whole system dynamics, which is also a starting point for revealing the mechanisms underlying the dynamics of agent-based complex systems (Grimm et al. 2005). This broadens the scope of existing concepts and methods, and even suggests the exciting possibility of a unified theory for apparently distinct financial and ecological complex systems.

## Authors’ contributions

YL designed the study and analyzed the data; YL, JR and VG performed the mathematical analyses and computer simulations; YL, JR and VG wrote the manuscript, with inputs from all authors. All authors gave final approval for publication.

## Acknowledgements

We are grateful to Stephen P. Hubbell for sharing the BCI dataset. The BCI forest dynamics research project was made possible by the U.S. National Science Foundation, numerous organizations and private individuals. We thank the field workers of the Center for Tropical Forest Science and the Smithsonian Tropical Research Institute for having made the BCI data sets publicly available. We thank Christophe Hurlin for his suggestions on panel data analysis and econometrics. We thank Robert Schlicht for his suggestions on statistical power. We also thank Jonathan Chase and Ryan Chisholm for their valuable comments.

## Note

This work and manuscript was finished on 27 April 2018 and was submitted to bioRxiv on 11 November 2021.

