## Appendix A for "Tropical forest dynamics correspond to fair games in economic theory of financial markets"

**symmetric, stabilizing, equalizing and neutral concepts**

1. German Centre for Integrative Biodiversity Research (iDiv) Halle-Jena-Leipzig, Puschstrasse 4, 04103 Leipzig, Germany; 2. Helmholtz Centre for Environmental Research − UFZ, Department of Ecological Modelling, Permoserstraße 15, 04318 Leipzig, Germany; 3. College of Life Sciences, Northwest University, 710069 Xi’an, P. R. China; 4. Division of Biology, Imperial College London, Silwood Park Campus, Ascot, Berkshire, SL5 7PY, UK; 5. Institute of Forest Growth and Computer Science, Technische Universität Dresden, P.O. 1117, 01735 Tharandt, Germany; 6. Institute of Biology / Geobotany and Botanical Garden, Martin Luther University Halle-Wittenberg, Am Kirchtor 1, 06108 Halle, Germany; 7. Max-Planck Institute for Biogeochemistry, Hans-Knöll-Str. 10, 07745 Jena, Germany

*These authors contributed equally to this work.

***Relationship to symmetric models***

The multiple species dynamic model in a community can be written as:

ln(*N_i,t_*) = *α_i_* + *β_i_* ln(*N_i,t_*_−1_) + *ε_i,t_* *i* = 1, 2, …, *S* ; *t* = 1, 2, 3, ... (A1)

where *N_i,t_* is the number of individuals in species *i* at time *t*. Note that this is a restatement of Equation (7) from the main text. In the context of this equation, a **symmetric model** is one where *α_i_* , *β_i_* and *ε_i,t_* are all independent of *i* so that species labels can be swapped without changing model dynamics. Symmetric models thus rule out the possibility of niches in a form where different species prefer different habitats or strategies.

***Stabilizing forces and equalizing forces***

An **equalizing mechanism** (Chesson 2000) is one where coexistence of species is promoted by individuals tending to have equal fitness, this is not the same as symmetry, which may include frequency dependence but rules out species that are different but nevertheless have equal fitness (e.g., because of a trade-off). To see the circumstances under which equalizing mechanisms are prevalent in our model we can rearrange Equation (A1) to give

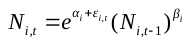
 (A2)

meaning that the average individual fitness is given by

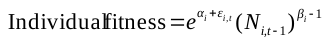
 (A3)

Where *α_i_* = 0 and *β_i_* = 1 in Equations (A1-A3) the system is a Martingale and there are strong equalizing forces; individual fitness is then given by $e^{\epsilon_{i,t}}$ where $\epsilon_{i,t}$ is stochastic and has mean 0. One can see this intuitively by considering the deterministic version of the equations where *ε_i,t_* = 0 for all *i*, *t* giving us *N_i,t_* = *N_i,t_*_−1_ and an individual fitness of unity in Equation (A3). The population growth rate for all species is thus independent of past abundances and is unpredictable (Main text Fig. 1f). As a result, the abundances of species do not necessarily return to their previous levels after stochastic events such as environmental disturbances (Main text Fig. 1e, f). In practice, coexistence under equalizing forces means birth and death rates are equal when averaged over time and individual fitness equals 1. In the more general case where we allow *α_i_* = *α* > 0 whilst *β_i_* = *β =* 1 the system is no longer a Martingale. There may still be strong equalizing forces at play, but individual fitness is given by exp(*α+ε_t_*) which on average is now > 1. Species will coexist still, but the system as a whole will grow in an unbounded way, in economic terms this means one cannot develop a winning *strategy* as such because everyone will win regardless. Conversely if *α_i_* = *α* < 0 whilst *β_i_* = *β =* 1 everyone will lose regardless and the system will collapse.

If fitness is not equalized among species in the community, then stable coexistence requires **stabilizing mechanisms**, which generate negative frequency-dependent regulations (Chesson 2000). In such cases, the population dynamics are not martingales and abundances of species will tend to return to previous levels after perturbations (e.g., environmental disturbance). Making the system in Equations (A1-A3) deterministic by setting *ε_i,t_* = 0 for all *i*, *t*, sheds light on this and also gives an indication of how the stochastic counterpart (where *ε_i,t_* is a distribution with zero mean) will behave. In the deterministic case, species *i* approaches an equilibrium abundance of
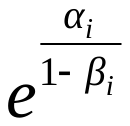
(found by setting $N_{i,t}=N_{i,t-1}$ in Equation A2 and rearranging). This could still be a strictly symmetric model as well, provided that *α_i_* = *α* and *β_i_* = *β* for all values of *i* so that all species approach the same equilibrium abundance. Deterministic models involving equalizing forces (*α_i_* = 0; *β_i_* = 1) can be constructed, but nothing interesting happens: species abundances simply don’t change and the equilibrium abundance equation
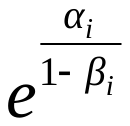
 is undefined because it depends instead on the unknown initial abundances.

***Restrictions on α and β***

Whilst the system given in Equation (A1) has well defined behavior for all values of its parameters *α_i_* and *β_i_* there are only certain values that make biological sense. For example $\left| \beta\right|\ll1$ means that the abundance of a species in the next time step is barely connected with its abundance in the present time step; this is not realistic for most biological systems. Similarly, away from *β* = 1 case of pure equalization, the system tends towards an abundance of
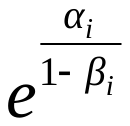
 which should not be too large nor too small and certainly not less than unity. Together these conditions place considerable restriction on the possible regions of parameter space likely to chosen as optimal by any fit of the model to empirical data. These regions are shown as white space in Figure A1.

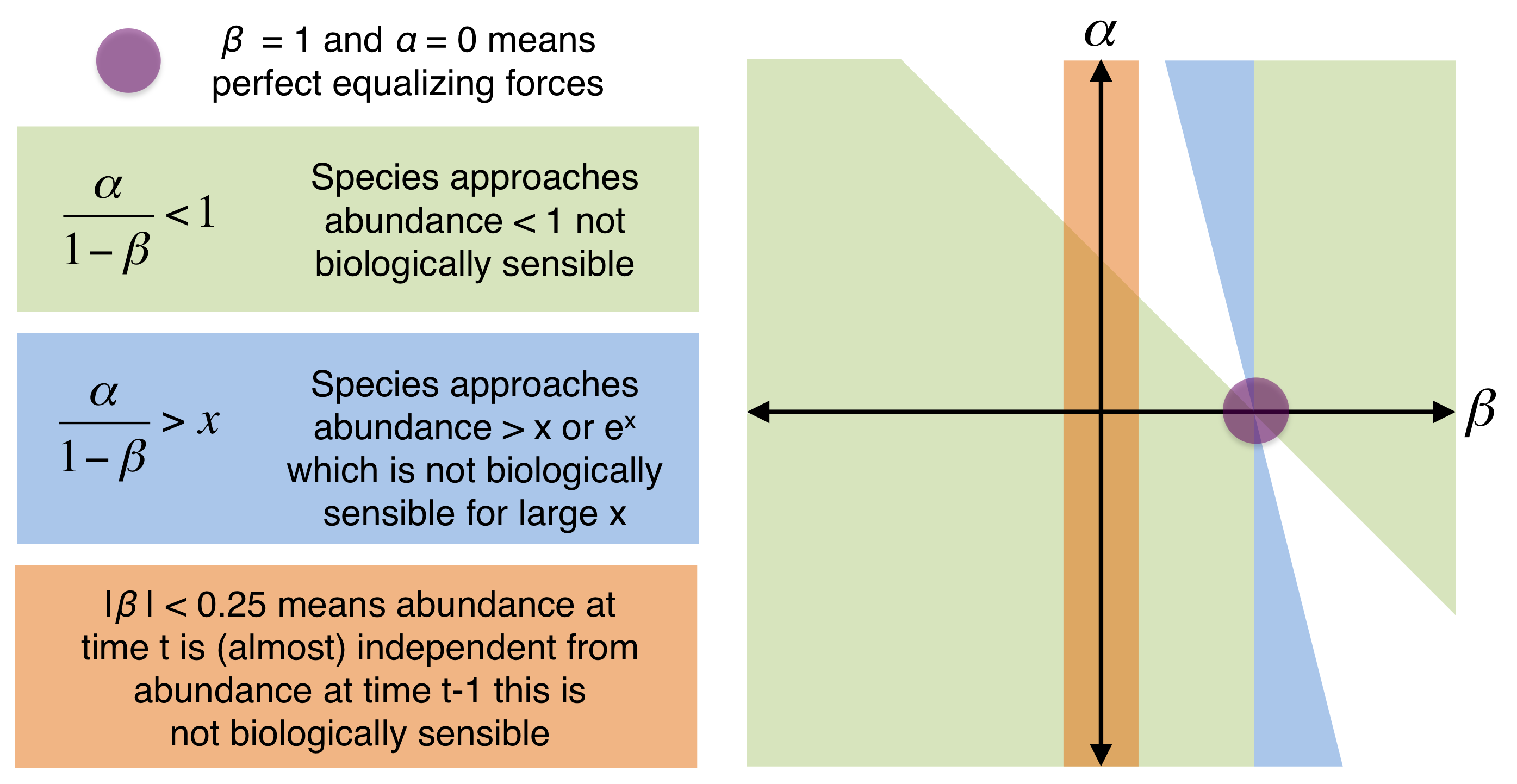

**Figure A1.** The parameter space for Equation (A1) based on varying *α* and *β*. The whitespace shows the only regions expected to correspond to biologically reasonable scenarios.

***Relationship to basic non-spatial neutral models***

If we now consider the model on arithmetic axes as described in Equation (6) of main text (*N_t_* = *α* + *β N_t_*_−1_ + *ε_t_*) we obtain a system that may be symmetric, stabilizing, equalizing or any combination of these based on the values of *α* and *β*; but where individuals rather than species are the focus. Such a model can relate to individual based ecological **neutral theory** (Hubbell 2001). Let us first consider the most basic non-spatial neutral model containing *J* individuals. In each model time step, an individual dies and is replaced by the offspring of another individual so the abundance *N_i,t_* of a species *i* at time *t* may increase by one, decrease by one, or remain constant compared to its former abundance *N_i,t-1_*. The probability of an increase in abundance is given by

$\frac{\left( J-N_{i,t-1} \right)}{J}\cdot\frac{\left( N_{i,t-1} \right)}{\left( J-1 \right)}$	(A4)

The first term giving the probability that the dead individual was not from species *i* and the second term giving the probability that the newborn is offspring of species *i*. The probability of a decrease in abundance can similarly be calculated and turns out to be identical to that of an increase. So in this case

*N_i,t_* = *N _i,t_*_−1_ + *ε_i,t_*	(A5)

Which is the same as Equation (6) with *α* = 0; *β* = 1 making species abundances a Martingale and indicating only equalizing forces. It is important to note that *ε_i,t_* has mean zero, but is a discrete distribution rather than a normally distributed one. If we consider that one time unit in Equation (A5) equates to a large number of time steps in the model then *ε_i,t_* can become normally distributed by the central limit theorem, but this is not guaranteed because *ε_i,t_* itself depends on *N_i,t_*. A further consideration is that zero is an absorbing state corresponding to species extinction, a consequence of studying Martingales in arithmetic (rather than logarithmic) space; this skews the *ε_i,t_* distribution away from a normal. The requirement for a martingale on arithmetic space to predict the same as a neutral model is therefore $1\ll N_{i,t}\ll J$so that small changes to *N_i,t_* do not significantly alter *ε_i,t_* and the absorbing state of $N_{i,t}=0$ is far away.

***Relationship to neutral models with speciation and/or immigration***

In classic neutral theory, species that go extinct are replaced in a dynamic equilibrium by new species that arrive either by speciation or by immigration from outside the community. We can deal with both of scenarios by the same approach. Let *m* represents the probability that a newborn individual is the offspring from an individual outside the system and let *S_i_* represents the probability that such an individual will be of species *i*. To model the case where all new individuals are new species we simply set $S_{i}=0$ for all values of *i*. The system can be described by the following set of equations.

$\begin{matrix} x=P\left( N_{i,t}=N_{i,t-1}+1 \right)=\frac{\left( J-N_{i,t-1} \right)}{J}\cdot\left( \frac{N_{i,t-1}\left( 1-m \right)}{\left( J-1 \right)}+mS_{i} \right) \\ y=P\left( N_{i,t}=N_{i,t-1}-1 \right)=\frac{N_{i,t-1}}{J}\cdot\left( \frac{\left( {J-N}_{i,t-1} \right)\left( 1-m \right)}{\left( J-1 \right)}+m\left( 1-S_{i} \right) \right) \\ z=P\left( N_{i,t}=N_{i,t-1} \right)=1-x-y \end{matrix}$	(A6)

The expected value of $N_{i,t}$ can be calculated as

$E\left( N_{i,t} \right)=x\cdot\left( N_{i,t-1}+1 \right)+y\cdot\left( N_{i,t-1}-1 \right)+ z\cdot N_{i,t-1}$	(A7)

By substituting the equations labeled (A6) into Equation (A7) we obtain

$E\left( N_{i,t} \right)=mS_{i}+N_{i,t-1}\left( 1-\frac{m}{J} \right)$	(A8)

Which can be directly related to Equation (6) in the main text with *α_i_* = $mS_{i}$and *β_i_* = $\left( 1-\frac{m}{J} \right)$. If we consider the classic case of a well-mixed metacommunity with speciation at rate $\nu$ and size *J*_M_, these equations reduce to *α_i_* = $0$and *β_i_* = $\left( 1-\frac{\nu}{J_{M}} \right)\approx1$ (because no immigration gives *S_i_* = 0, low speciation rate gives $\nu\ll1$ and large size gives *J*_M_ $\gg1$).

The stabilizing and/or equalizing forces at work maintaining local species diversity may well be different to those maintaining global species diversity. If we now consider only the local community part of the spatially implicit neutral model (with a fixed external metacommunity), this need not be symmetric unless $S_{i}$ happens to be independent of *i*. Furthermore, it contains stabilizing forces as well as equalizing ones (Chisholm et al. 2014) which can be seen in our equations because *β_i_* = $\left( 1-\frac{m}{J} \right)$≠1. One special case is *m* = 0 which reduces the local community case to that of a small isolated metacommunity with no speciation, something that is symmetric and possessing equalizing forces only.

In the deterministic equivalent of the arithmetic scale martingale shown in Equation (6), species *i* approaches an equilibrium abundance of $\frac{\alpha_{i}}{1-\beta_{i}}$. This has similar properties to the counterpart on logarithmic scales shown earlier, in particular that it is undefined when *α_i_* = 0 and *β_i_* = 1. If we substitute in the values for *α_i_* and *β_i_* in terms of neutral theory parameters $m, S_{i}$and *J* we obtain $\frac{mS_{i}}{1-\left( 1-\frac{m}{J} \right)}=JS_{i}$ which makes intuitive sense as results in the local community being divided up between species proportional to their outside abundances and immigration pressure given by $S_{i}$. Similar restrictions on possible values of *α* and *β* apply as shown in Figure (A1), however the conditions are slightly less strict since $\frac{\alpha_{i}}{1-\beta_{i}}$ has not been exponentiated.

***Summary***

- Symmetric models in the context of Equations (6) and (7) of the main text are ones where the values of *α_i_* , *β_i_* and *ε_i,t_* are all independent of *i*, which is the species label.
- On logarithmic axes, a helpful deterministic analogy to Equation (7) gives the result that each species *i* approaches an equilibrium abundance of
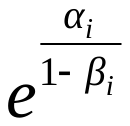
.
- Where *α_i_* = 0 and *β_i_* = 1 we have strong equalizing forces in the model, the time averaged net fitness of an individual in the model is unity, and equilibrium abundance is undefined.
- Where *α_i_* ≠ 0 and *β_i_* = 1 we still have strong equalizing forces. However, the system as a whole will either explode without bounds (*α_i_* > 0) or collapse (*α_i_* < 0).
- If *β_i_* ≠ 1 then we have a stabilizing component to the system and there is an inclination for species to head towards an equilibrium abundance; this inclination may be strong or weak depending on the *ε_i,t_* and the other model parameters.
- Forces maintaining global diversity may be different from those maintaining local diversity; such forces may be stabilizing and/or equalizing.
- The assumption of ecological equivalence in the original neutral model (Hubbell 2001) means that demographic rates of an individual are independent of species identity. This in turn makes the resulting model a symmetric model, but a symmetric model need not be neutral and may include stabilizing and/or equalizing forces.
- The metacommunity part of classic neutral theory (Hubbell 2001) is non spatial and with some background speciation at a low rate; this corresponds to *α_i_* = 0 , *β_i_* = $\left( 1-\frac{\nu}{J_{M}} \right)\approx1$. It thus corresponds to an equalizing mechanism analogous to the financial dynamics asserted by ‘efficient market theory’, only on arithmetic rather than logarithmic space.
- Neutral Theory only corresponds to efficient market theory; the two are not identical because of differences in the stochastic elements of the two systems. For abundant species in an even larger system, however, the differences are expected to become negligible and perfect equivalence should emerge.
- If one considers just the local community part of the classic neutral model separate from the metacommunity part, it may not be symmetric, and furthermore dispersal may act as a stabilizing force. Community size *J*, dispersal limitation *m* and metacommunity abundances *S_i_* can be used to determine equivalent values for to *α_i_* and *β_i_* in Equation (6) using *α_i_* = $mS_{i}$and *β_i_* = $\left( 1-\frac{m}{J} \right)$.
- In the deterministic equivalent of arithmetic scale martingale shown in Equation (6), species *i* approaches an equilibrium abundance of $\frac{\alpha_{i}}{1-\beta_{i}}$ which in the equivalent neutral model is given by $JS_{i}$.
- In both arithmetic and logarithmic space there are restrictions on the possible values for *α_i_* and *β_i_* that would be considered biologically reasonable. The restrictions are more severe in the case of logarithmic space as a small change in parameters can result in large changes in real abundances.

***References***

Chesson, P. 2000. Mechanisms of maintenance of species diversity. Annual Review of Ecology and Systematics 31:343–366.

Chisholm, R. A., R. Condit, K. A. Rahman, P. J. Baker, S. Bunyavejchewin, Y.-Y. Chen, G. Chuyong, et al. 2014. Temporal variability of forest communities: empirical estimates of population change in 4000 tree species. Ecology Letters 17:855–865.

Hubbell, S. P. 2001. The Unified Neutral Theory of Biodiversity and Biogeography. Princeton University Press.
