## Appendix B for "Tropical forest dynamics correspond to fair games in economic theory of financial markets"

1. German Centre for Integrative Biodiversity Research (iDiv) Halle-Jena-Leipzig, Puschstrasse 4, 04103 Leipzig, Germany; 2. Helmholtz Centre for Environmental Research − UFZ, Department of Ecological Modelling, Permoserstraße 15, 04318 Leipzig, Germany; 3. College of Life Sciences, Northwest University, 710069 Xi’an, P. R. China; 4. Division of Biology, Imperial College London, Silwood Park Campus, Ascot, Berkshire, SL5 7PY, UK; 5. Institute of Forest Growth and Computer Science, Technische Universität Dresden, P.O. 1117, 01735 Tharandt, Germany; 6. Institute of Biology / Geobotany and Botanical Garden, Martin Luther University Halle-Wittenberg, Am Kirchtor 1, 06108 Halle, Germany; 7. Max-Planck Institute for Biogeochemistry, Hans-Knöll-Str. 10, 07745 Jena, Germany

*These authors contributed equally to this work.

**Population dynamics and ecological martingales**

In probability theory, a ‘martingale’ is a stochastic process where the next expected value in the time-series is equal to the previous value, and past information never helps to reduce the uncertainty about future outcomes. A martingale reflects the outcome of a ‘fair game’ model because it excludes the possibility to use a winning strategy that is based on past game history for improving expected future outcomes (Fama 1970; Barnett and Serletis 2000; Poitras 2010).

Symbolically, a discrete-time martingale, with respect to certain conditions, is defined as

*E*(*Y_t_* | *X*_0_, …, *X_t_*_−1_) = *Y_t_*_−1_	*E*(|*Y_t_*|) < ∞; *t* = 1, 2, ..., *T*	(B1)

where *E* is the expected value, *Y_t_* the variable of interest observed at time *t*, and *X_t_* represents the system’s condition given at time *t*. Equation (B1) says that the expected value of the next observation *Y_t_*, given the sequence of previous conditions, *X*_0_, …, *X_t_*_−1_, is equal to the last observation, *Y_t_*_−1_.

To check whether the martingale model can be applied to ecological communities, we describe the expected population size of each species with respect to their ecological fitness in a general way as:

*E*(*N_i,t_* | Θ*_i,_*_0_, ..., Θ*_i,t_*_−1_) = *E*(*λ_i,t_* | Θ*_i,_*_0_, ..., Θ*_i,t_*_−1_) *N_i,t_* _−1_	*i* = 1, 2, ..., *S* ; *t* = 1, 2, ..., *T*	(B2)

where *N_i,t_* is the abundance of species *i* at time *t*; *λ_i,t_* is the finite population growth rate of species *i* at time *t*; and sequence {Θ*_i,t_*} stands for the ecological fitness of species *i*.

A reasonable estimation of fitness can be based on the ratio *R_i_*_,_*_t_* of birth (*b_i_*) and death (*d_i_*) rate (Hubbell 2001, 2008; Chave 2004; Allouche and Kadmon 2009; Lin et al. 2009; Zhang et al. 2012), which gives:

Θ*_i_*_,_*_t_* = *R_i_*_,_*_t_* = *b_i_*_,_*_t_* / *d_i_*_,t_	*i* = 1, 2, ..., *S*; *t* = 1, 2, ..., *T*	(B3)

If fitness equalizing processes hold according to demographic trade-offs, so that a higher birth rate is equalized by a higher death rate, then *R_i_*_,_*_t_* is constant and the same for all co-occurrence species (Parsons and Quince 2007; Allouche and Kadmon 2009; Lin et al. 2009; Ostling 2012; Zhang et al. 2012). A community where fitness is equalized among all species, however, is at demographic equilibrium (*R_i_* = *b_i_* / *d_i_* = 1) for all *i* (Chave 2004; Ostling 2012). According to the community models of Lin et al. (2009), Ostling (2012) and Zhang et al. (2012), when speciation is absent, the expected species’ abundances should neither increased nor decreased, which leads to

*E*(*N_i,t_* | *R_i,_*_0_, ..., *R_i,t_*_−1_) = *N_i,t_* _−1_	*i* = 1, 2, ..., *S* ; *t* = 1, 2, ..., *T*	(B4)

Equation (B4) indicates that with respect to fitness equalizing, the species’ abundance should show no consistent trends of change, and the abundance time-series of all *S* species in the community are martingales. Note that the real birth rate is slightly smaller than the death rate when there is speciation because a species can also be replaced by a new species. On the timescales we are evaluating, however, speciation is insignificant and therefore we only consider dynamics without speciation. Immigration is also a consideration in our model, we implicitly assume immigration-emigration equilibrium in our model for each species. We discuss in Appendix A the interpretation of immigration as a stabilizing force that is consequently incorporated into our analyses implicitly. Furthermore, previous work suggests that immigrants into the forest plot on Barro Colorado Island contribute less to the recruitment (Chisholm and Lichstein 2009).

According to Equations (B2), (B3) and (B4), we have

*E*(*λ_i,t_* | *R_i,_*_0_, ..., *R_i,t_*_−1_) = 1	*i* = 1, 2, ..., *S* ; *t* = 1, 2, ..., *T*	(B5)

which indicates equalized fitness. Population growth is multiplicative, so the distribution of the finite population growth rate of a species through time is approximately log-normal (Condit et al. 1999; Chave 2004; Hubbell 2008; Chisholm et al. 2014). Consequently, the general form of population dynamics, on a logarithmic scale, under ecological equalizing conditions should follow:

ln(*N_i,t_*) = ln(*N_i,t_*_−1_) + *ε_i,t_*	*i* = 1, 2, ..., *S* ; *t* = 1, 2, ..., *T*	(B6)

where *ε_i,t_* is known as the martingale difference that represents a stochastic process with the property of zero mean and independent distribution. Equation (B6) is similar to the Gaussian random walk model, which is also widely used in economics for describing financial dynamics. Of particular importance is that in Gaussian random walk model, the stochastic term represents independent draws from a normal distribution with mean zero and variance *σ_i_*^2^ (white noise). Consequently, the Gaussian random walk is a more restrictive case of martingale process (Barnett and Serletis 2000). Here, for convenience of analysis, we employed this restrictive assumption of martingale process where *ε_i,t_* stands for a white noise process. Equation (B6) can be rewritten as

ln*λ_i,t_* = ln(*N_i,t_*) – ln(*N_i,t_*_−1_) = *ε_i,t_*	*i* = 1, 2, ..., *S* ; *t* = 1, 2, ..., *T*	(B7)

which suggests that under equalizing mechanisms, for each species in the community, the time-series of population growth rate is uncorrelated and follows a mean zero normal distribution (white noise).

**Unit root tests with panel data**

The stochastic process represented by Equation (B6) is non-stationary (its mean and variance are changing over time), but the series of differences between consecutive values (Equation B7) is stationary (its mean and variance are time independent). A stochastic process with this property is referred to as ‘unit root’ process (Dickey and Fuller 1981; Barnett and Serletis 2000). A martingale implies a unit root process. In economics, unit root tests are widely used to test the martingale hypothesis for single time-series (Barnett and Serletis 2000; Poitras 2010). The general model for testing unit root process is

*Y_i,t_* = *α_i_* + *β_i_Y_i,t_*_−1_ + *ε_i,t_*	(B8)

where *α_i_* and *β_i_* are constants. When *Y_i,t_* denotes ln(*N_i,t_*), the log-transformed abundance of species *i* at time *t*, equation (B8) is identical to the linear form of the stochastic Gompertz model. The Gompertz model is frequently used in ecological studies for modelling single population dynamics and for testing negative density-dependence. It is flexible enough to represent both variations among different species and the demographic and environmental stochasticity (Ives et al. 2003; Dennis et al. 2006). On a logarithmic scale, the Gompertz model is a linear first-order autoregressive time-series model:

ln(*N_i,t_*) = *α_i_* + *β_i_* ln(*N_i,t_*_−1_) + *ε_i,t_*	(B9)

Also, we can rewrite Equation (B9) as

ln*λ_i,t_* = ln(*N_i,t_*) − ln(*N_i,t_*_−1_) = *α_i_* + (*β_i_* – 1) ln(*N_i,t_*_−1_) + *ε_i,t_*	(B10)

where *α_i_* and *β_i_* are species-specific constants that can represent variations among species in population growth rate and different responses to environmental factors. When *α_i_* = 0 and *β_i_* = 1, equation (B9) is the same as Equation (B6), which says that the stochastic process has a unit root, in agreement with the martingale process of equalizing mechanisms; and Equation (B10) is identical to Equation (B7), which implies that the species growth rate is independent of species abundance. If *α_i_* ≠ 0 and |*β_i_*| < 1 (|*β_i_*| >> 1 is biologically unlikely, Ives *et al.* 2003; Dennis *et al.* 2006), the dynamics of ln(*N_i,t_*) approach a stationary distribution with a mean of *α_i_* /(1− *β_i_*) and a variance of *σ_i_*^2^/(1− *β_i_*^2^). This dynamical process does not have a unit root, and hence is not a martingale process, and in ecological terms, such a dynamical process implies strong negative frequency-dependent regulation of stabilizing mechanisms (Ives et al. 2003). Analytically, the coefficient *β_i_* in Equations (B9) and (B10) can be used to measure the strength of frequency-dependent regulation, which ranges from 0 (strict frequency-dependent regulation) to 1 (no frequency-dependent regulation) (Ives et al. 2003; Dennis et al. 2006). Some studies suggest that when *β_i_* is very close to 1, the ordinary tests on single time-series often fail to distinguish unit root null (equalizing mechanisms) from the alternative hypothesis (stabilizing mechanisms) (Dickey and Fuller 1981; Ives et al. 2003). Therefore, we employ panel data tests that have more power than ordinary tests (Hsiao 2003, 2007). A *β_i_* very close to 1 indicates a weak stabilizing mechanism that would be in any case less ‘critical’ to detect.

The Augmented Dickey-Fuller (ADF) test (Dickey and Fuller 1981) is widely used for testing unit root process in a single time-series. The ADF test is applied to the model

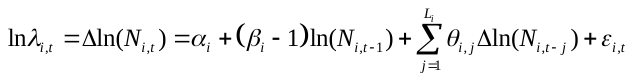
 (B11)

where *L_i_* the number of augmenting lags, is introduced in order to reduce autocorrelation of error terms, and is determined by minimizing the Akaike information criterion or minimizing the Schwartz Bayesian information criterion, we have *L_i_* in our tests determined automatically by using EViews 7 (http://www.eviews.com/ ). Then ordinary least squares and *t*-statistic can be applied for testing *α_i_* = 0 and *β_i_* =1, as for testing unit root process in single time-series.

Because we are going to evaluate multiple time-series of co-occurring species in a community simultaneously, and the observed time-series are relatively short, we cannot use ordinary unit root tests (e.g., ADF test) that are only applicable to single and long time-series. We therefore employ the newly developed techniques of unit root tests with panel data, which was widely used for analyzing survey data consisting of many short but associated time-series (often more than dozens of time-series with only five or six time points) (Hsiao 2003, 2007). The basic concept of panel unit root tests is similar to that of meta-analysis: it combines the statistical information of all the time-series across different sections (species in our case) to draw an overall conclusion (Maddala and Wu 1999; Hsiao 2003). To make the results of our panel unit root test more robust, we applied three different established methods of panel unit root test in this study: the IPS test (Im et al. 2003), ADF-Choi Z test (Choi 2001) and ADF-Fisher χ^2^ test (Maddala and Wu 1999). All three panel tests are based on ADF test. The IPS test is based on the average of *t*-statistics of ADF test for all time-series:

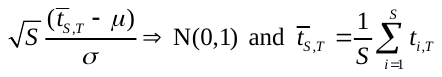
 (B12)

where *t_i,T_* (*i* = 1, 2, ..., *S*) denotes the *t*-statistics for testing unit roots, N(0,1) denotes the standard normal distribution, and *E*(*t_i,T_*)=*μ* and *V*(*t_i,T_*)=*σ* are the mean and variance, respectively. The ADF-Choi Z test and ADF-Fisher χ^2^ test are both Fisher-type tests that are based on combining the significance levels (*P*-values) of the ADF test for each time-series, but with a minor difference in the underlying algorithms (Maddala and Wu 1999; Choi 2001). The ADF-Choi Z test is defined as

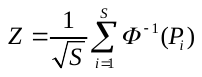
 (B13)

where *Φ*(·) is the standard normal cumulative distribution function, and *P*_i_ is the *P*-value of ADF test for *i*. The ADF-Fisher χ^2^ test is defined as

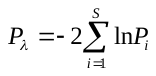
 (B14)

which has a χ^2^ distribution with 2*S* degrees of freedom. Simulation studies have demonstrated that the panel unit root tests we used here are much more powerful than single unit root tests (Maddala and Wu 1999; Hsiao 2003, 2007; Im et al. 2003; Hlouskova and Wagner 2006), and the ADF-Fisher χ^2^ test is also applied when there are cross-sectional correlations (Maddala and Wu 1999). In addition, the tests we used here allow unbalanced panels (Maddala and Wu 1999). In an unbalanced panel, the number of time periods is not the same for all individual time-series which assumes unequally spaced time steps.

We ran all three tests, using the Gompertz model (Equation B9) as the underlying model. According to the statistical hypothesis of panel unit root tests used here, our null- and alternative hypotheses are defined as:

*H_0_*: *α_i_* = 0 and *β_i_* = 1, *i* ∀ *S* (Equalizing mechanisms only)

*H_A_*: *α_i_* ∈ **R** and *β_i_* < 1, *i* ⊆ *S* (Stabilizing mechanism components)

The three panel tests we used here are also used more widely and are relatively flexible compared to other panel tests. According to the statistical hypothesis of the panel unit root tests used here, our null hypotheses means for all species that their time-series are martingales corresponding to equalizing mechanisms; whilst our alternative hypothesis means some (but not necessarily all) species have time-series that are not martingales. The alternative hypothesis of the three tests we used here also considered the possibility of a nested structure where stabilizing mechanisms applied only to some subset of species. This is different from other panel unit root tests such as the LLC test (Levin et al. 2002), in which the alternative hypothesis was *H_A_*: *α_i_* = *α* ∈ **R** and *β_i_* = *β* < 1, and all time-series share common coefficients.

**Panel data from a species-rich tropical forest**

We used a species abundance data set of the 50 ha-plot on Barro Colorado Island (BCI), Panama (Condit 1998; Condit et al. 1999). Since 1981, seven censuses were accomplished (1981, 1985, 1990, 1995, 2000, 2005 and 2010), all free-standing trees in the plot with stem diameter ≥ 1 cm (at 1.3 m above the ground) were mapped, tagged, and identified to the species level. The census data were collected according to standard methods of the CTFS studies (Condit 1998; Condit et al. 1999). In contrast to most earlier analyses of this unique data set, we used census data of each species for panel analysis. Thus, the panel data of the BCI plot contains 322 species (sections) and seven census time points with a census interval of 5-years (the interval between first census at 1981 and second one was four years but, to simplify, we treat this as a 5-year interval in our analysis).

We applied our analysis to the full data set, and separately to saplings (1 cm ≤ stem diameter < 10 cm) and to large trees (stem diameter ≥ 10 cm) only. Several species were excluded from our analysis as their time-series were too short or were not applicable (Supplementary data). Panel data analyses were performed in EViews 7 (http://www.eviews.com/), RVN test (package ‘lawstat’) and Ljung-Box test were conducted using R 2.15.3 (http://www.r-project.org/), BCI data set were processed using CTFS R package (http://ctfs.arnarb.harvard.edu/Public/CTFSRPackage/).

**Individual-based simulation model details**

To evaluate the power of unit root tests, we also developed an individual-based tree competition model. In this simulation model, species coexist and compete for 10,000 empty grid-cells and species dynamics can be switched to either stabilizing or equalizing mechanisms. Our individual-based demonstration model of multiple tree species dynamics could mechanistically produce both stationary processes based on stabilizing mechanisms, and non-stationary processes based on equalizing mechanism (martingales). Species dynamics in our simulation model only contain birth-death processes. Under the stabilizing mechanism, species growth rates are assigned in an orderly manner; species with lower intrinsic birth rates are assigned a higher competitive ability (priority of grid occupation), this trade-off between population growth rate and competitive ability promotes a stable coexistence among those species. Under the equalizing mechanism, we assumed all species have equal overall fitness based on demographic trade-offs. The overall fitness given by the ratio of intrinsic birth/death rate (net reproductive rate) is equal to 1.5 for all species. Whilst in Hubbell’s (2001) neutral models this ratio was set to 1.0 as a simplification, a value greater than 1 is more realistic in neutral models (Ostling 2012). We also used a ratio value 1.0 in our model for comparison and found that this did not change our results or conclusions because the dynamical processes produced by those models are still martingales. In both mechanisms, the intrinsic birth rates of species ranged from 0.078 to 0.822 so as to represent a varied characteristic of species demography. The total replacement of individuals in our simulation model is set to 2% per time step which is similar as in the BCI forest (about 2% per year). We introduced stochasticity to individual vital rates and establishment rate on empty grid-cells, so that the simulated tree species dynamics under different mechanisms also contains temporal stochasticity such as demographic and environmental stochasticity (Fig. S7). For each simulation, we run 1030 time steps and only use 7 time points (time step 1000, 1005, 1010, 1015, 1020, 1025 and 1030) for panel unit root tests, so as to reproduce a dataset that is similar as the empirical one in the in the BCI forest. We repeat 10 times for each simulation by changing the seed of pseudo-random number generator. Our simulation model and experiments were implemented in a free simulation platform NetLogo 4.1.3 (Appendix C, model interface, source code and simulation experiments are contained in a .nlogo file, Wilensky 1999).
