## Supplemental Fig. S1-S6, Table S1-S3 for "Tropical forest dynamics correspond to fair games in economic theory of financial markets"

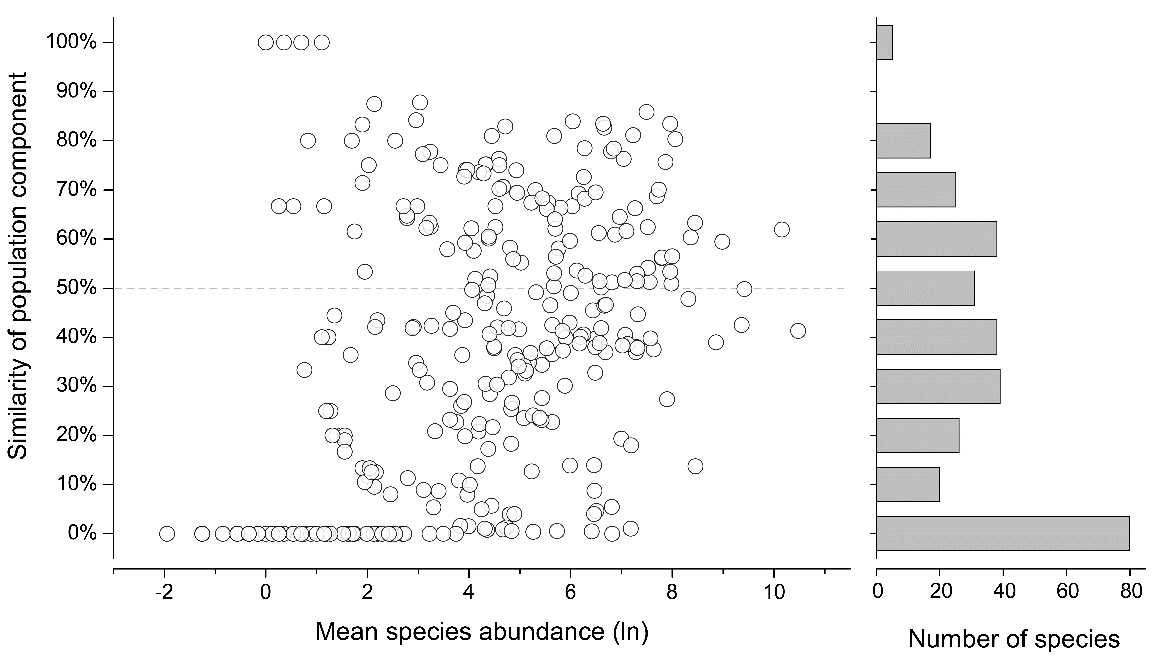

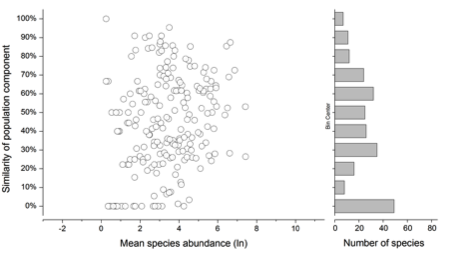

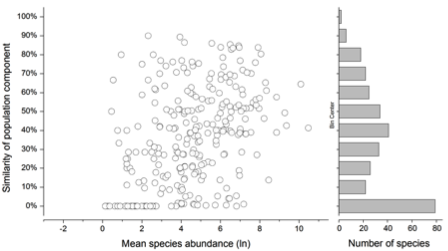

(a)

(b)

(c)

**Figure S1:** Temporal turnover (similarity of population component) over 29 years against mean abundance for tree species in a 50 ha-plot of species-rich tropical forest on Barro Colorado Island (BCI). Turnover was calculated with the Sørensen index given by the equation $\frac{2\cdot N_{s}}{\left( N_{0}+N_{f} \right)}\cdot100\%$ where *N*_s_ gives number of individuals that exist in both first and last census, *N*_0_, gives the number of individuals in the first census and *N*_f_ gives the number of individuals in the last census. All free-standing individuals with DBH ≥ 1 cm were included in the calculations. For species where Sørensen index could be calculated, more than 60% of species had a similarity less than 50%. (*a*) pooled data of all individuals. (*b*) data of all trees (DBH ≥ 10 cm). (*c*) data of all saplings (1 cm ≤ DBH < 10 cm).

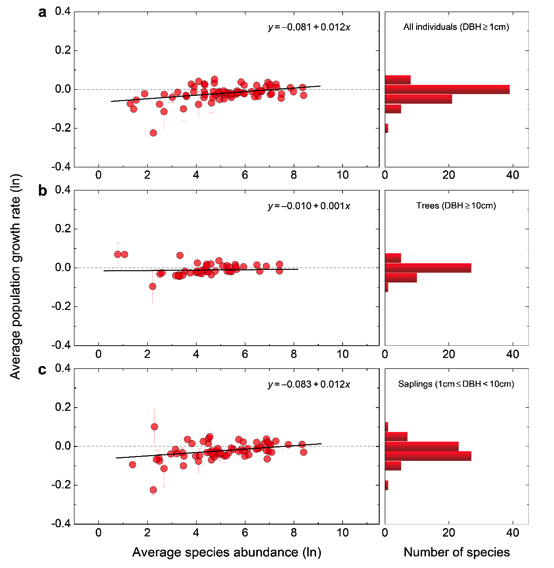

**Figure S2:** Population growth rate (mean ± 1 SD) distribution of only those tree species that differ from ‘fair game’ dynamics of zero in a 50 ha-plot of species-rich tropical forest on Barro Colorado Island. The solid lines show fitted ordinary least-squares regression. (*a*) pooled data of all individuals (74 species, *r*^2^ = 0.224, *P* < 0.001). (*b*) data of all trees (43 species, *r*^2^ = 0.002, *P* = 0.809). (*c*) data of all saplings (64 species, *r*^2^ = 0.164, *P* < 0.001).

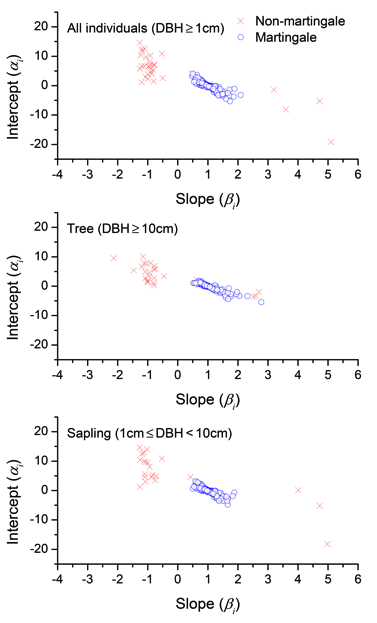

**Figure S3:** Estimation of species-specific parameters of abundance dynamics, *α_i_* and *β_i_*, by using the auto-regressive equation ln(*N_i,t_*) = *α_i_* + *β_i_* ln(*N_i,t_*_−1_). Standard major axis regression (type II model) was used for fitting, which is symmetric and assumes error in both variables. Blue circles: species whose 95% confidence interval of regression line is consistent with martingale properties. Red crosses: species whose 95% confidence interval of regression line is not consistent with martingale properties (37 species out of 276 species for all individuals; 32 species out of 203 species for trees; 30 species out of 257 species for saplings). When considering the distribution of parameter pairs *α_i_* and *β_i_* for each species *i*, we found patterns where certain areas of parameter space are not occupied by any species in Figure S4. These unoccupied regions usually correspond to solutions to the equations that are stabilized, but with unrealistically high or unrealistically low target abundances. The blank area of parameter space around *|β| ≈* 0 corresponds to an unrealistic case where species abundances in the next point of the time series are totally decoupled from the previous point in time (electronic supplementary material, Appendix A, Equation 7). Our tests cannot reject either the symmetric, equalizing (but not neutral) case where species abundances follow fair game dynamics on logarithmic space, or the symmetric and neutral case where abundances correspond to fair games on arithmetic space. There is marginally more support for the non-neutral case, however it is hard to distinguish between logarithmic and arithmetic scales from data where fluctuations in abundance are small in general.

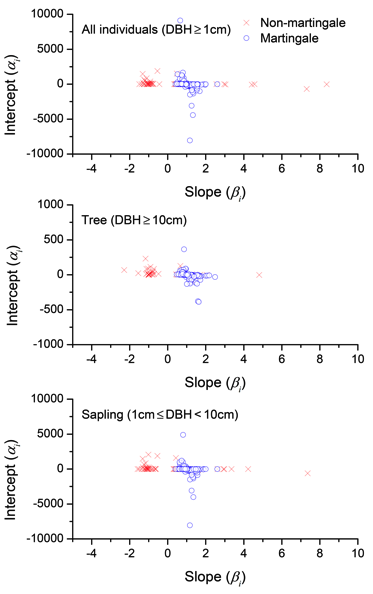

**Figure S4:** Estimation of species specific parameters of abundance dynamics, *α_i_* and *β_i_*, by using the auto-regressive equation *N_i,t_* = *α_i_* + *β_i_* *N_i,t_*_−1_ (Equivalent to Figure S3 but on arithmetic axes). A standard major axis regression (type II model) was used for fitting, this is symmetric and assumes error in both variables. Blue circles: species whose 95% confidence interval of regression line is consistent with martingale properties. Red crosses: species whose 95% confidence interval of regression line is not consistent with martingale properties (62 species out of 311 species for all individuals; 41 species out of 234 species for trees; 63 species out of 307 species for saplings).

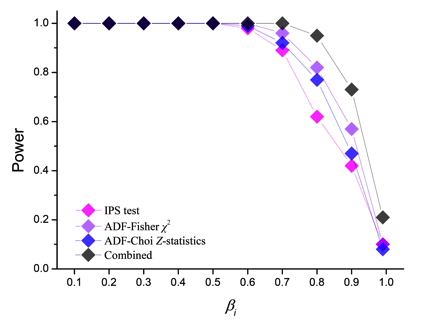

**Figure S5:** Statistical power of panel unit root tests on rejecting the martingale null hypothesis as a function of the strength (from 0.1 to 0.99) of stabilizing processes (arithmetic space). Note that the simulated panel data sets (each contains 322 time-series of 7 data points) were generated based on empirical data set of BCI. This meant that not all 322 species could be tested because some of the time-series are too short or incomplete. This graph accompanies Figure 4 of the main text, which shows the same results for a logarithmic space martingale model and panel unit root test.
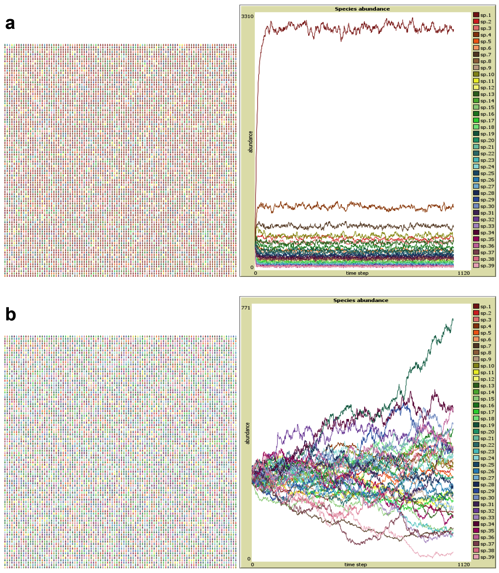

**Figure S6:** Snapshots of simulations of tree species dynamics. (*a*) dynamics based on a strong stabilizing mechanism. (*b*) dynamics based on only equalizing mechanisms. In the simulation model, time-series data sets of 39 species with 7 time points (time step 1000, 1005, 1010, 1015, 1020, 1025 and 1030) were used for testing the power of panel unit root tests. See Appendix B for full model details.
**Table S1.** Results of panel unit root tests for tree species abundance (arithmetic space) dynamics in a 50 ha-plot of species-rich tropical forest on Barro Colorado Island

|  | IPS test  Statistic (Probability) | ADF-Choi *Z*-statistics  Statistic (Probability) | ADF-Fisher *χ*^2^  Statistic (Probability) |
| --- | --- | --- | --- |
| All individuals (DBH ≥ 1 cm) | 0.513 (*P* = 0.696) | NA | 693.612 (*P* = 0.045) |
| Trees (DBH ≥ 10 cm) | 3.814 (*P* = 0.999) | NA | 432.129 (*P* = 0.972) |
| Saplings (1 cm ≤ DBH < 10 cm) | 1.047 (*P* = 0.852) | 2.966 (*P* = 0.999) | 684.027 (*P* = 0.038) |

Null hypothesis: the existence of unit roots for all species and non-stationary dynamics (martingale processes) predicted by equalizing mechanisms (see Appendix A for more details). Alternative hypothesis: for some (but not necessarily all) species the unit root does not exist meaning they are governed by the stationary dynamics predicted by stabilizing (niche-based) mechanisms. NA indicates a test statistic that is not applicable to the data set. Note it is not applicable to use the tests for all 322 species because some of the time-series are too short or incomplete (see detailed species lists in Appendix D).

**Table S2** Results of the tests for independence and randomness of tree species population growth rates in a 50 ha-plot of species-rich tropical forest on Barro Colorado Island.

|  | Number of species tested | Ljung-Box test  (Null: independence) | RVN test  (Null: randomness) |
| --- | --- | --- | --- |
| All individuals (DBH ≥ 1 cm) | 292 | 100% (*P* > 0.050) | 96.92% (*P* > 0.059) |
| Trees (DBH ≥ 10 cm) | 216 | 96.76% (*P* > 0.053) | 98.61% (*P* > 0.050) |
| Saplings (1 cm ≤ DBH < 10 cm) | 279 | 99.28% (*P* > 0.058) | 96.42% (*P* > 0.059) |

It is not appropriate to use the tests for all 322 species because some of the time-series are too short or incomplete. The percentages indicate the proportion of species for which growth rates are not significantly different from the null expectation of no autocorrelation.

**Table S3** Results of panel unit root tests for simulations of tree species dynamics based on stabilizing or equalizing mechanisms

|  | Random seed | IPS test  Statistic (Probability) | ADF-Choi *Z*-statistics  Statistic (Probability) | ADF-Fisher *χ*^2^  Statistic (Probability) |
| --- | --- | --- | --- | --- |
| Stabilizing mechanism | 0 | -3.333 (*P* = 0.000) | -4.430 (*P* = 0.000) | 133.688 (*P* = 0.000) |
|  | 1 | -2.172 (*P* = 0.015) | -3.284 (*P* = 0.001) | 116.102 (*P* = 0.003) |
|  | 2 | -2.147 (*P* = 0.016) | -2.639 (*P* = 0.004) | 111.903 (*P* = 0.007) |
|  | 3 | -2.879 (*P* = 0.002) | -4.303 (*P* = 0.000) | 125.391 (*P* = 0.001) |
|  | 4 | -3.327 (*P* = 0.000) | -4.505 (*P* = 0.000) | 134.917 (*P* = 0.000) |
|  | 5 | -2.557 (*P* = 0.005) | -3.835 (*P* = 0.000) | 120.803 (*P* = 0.001) |
|  | 6 | -2.725 (*P* = 0.003) | -4.389 (*P* = 0.000) | 122.756 (*P* = 0.001) |
|  | 7 | -4.141 (*P* = 0.000) | -4.713 (*P* = 0.000) | 146.969 (*P* = 0.000) |
|  | 8 | -2.593 (*P* = 0.005) | -3.670 (*P* = 0.000) | 119.423 (*P* = 0.002) |
|  | 9 | -4.258 (*P* = 0.000) | -5.428 (*P* = 0.000) | 151.813 (*P* = 0.000) |
| Equalizing mechanism | 0 | 0.429 (*P* = 0.666) | 0.158 (*P* = 0.563) | 67.314 (*P* = 0.800) |
|  | 1 | -0.533 (*P* = 0.297) | -0.917 (*P* = 0.180) | 91.563 (*P* = 0.140) |
|  | 2 | 0.217 (*P* = 0.586) | 0.053 (*P* = 0.521) | 83.376 (*P* = 0.318) |
|  | 3 | -1.172 (*P* = 0.121) | -1.143 (*P* = 0.127) | 100.830 (*P* = 0.042) |
|  | 4 | 0.159 (*P* = 0.563) | -0.123 (*P* = 0.451) | 76.771 (*P* = 0.518) |
|  | 5 | -0.194 (*P* = 0.423) | -0.498 (*P* = 0.309) | 81.689 (*P* = 0.365) |
|  | 6 | 0.579 (*P* = 0.719) | 0.263 (*P* = 0.604) | 82.524 (*P* = 0.341) |
|  | 7 | 0.187 (*P* = 0.574) | -0.036 (*P* = 0.486) | 82.911 (*P* = 0.331) |
|  | 8 | 1.165 (*P* = 0.878) | 1.290 (*P* = 0.902) | 67.182 (*P* = 0.804) |
|  | 9 | 0.082 (*P* = 0.533) | -0.208 (*P* = 0.418) | 82.982 (*P* = 0.329) |

Null hypothesis: the existence of unit roots for all co-occurrence species; non-stationary dynamics (martingale processes) governed by equalizing mechanisms. Alternative hypothesis: some (but not necessarily all) species the unit root does not exist meaning they are governed by the stationary dynamics predicted by stabilizing mechanisms. For each simulation, we repeat 10 times by changing the seed of pseudo-random number generator. All panel tests show powerful performances in identifying the underlying mechanisms that were based on the simulation results of 39 species with 7 time points (time step 1000, 1005, 1010, 1015, 1020, 1025 and 1030). For further details on the model see Appendix [fill in ] and Figure S7.
